# Melatonin suppression does not automatically alter sleepiness, vigilance, sensory processing, or sleep

**DOI:** 10.1101/2022.04.12.488023

**Authors:** Christine Blume, Maria Niedernhuber, Manuel Spitschan, Helen C. Slawik, Martin P. Meyer, Tristan A. Bekinschtein, Christian Cajochen

## Abstract

Pre-sleep exposure to short-wavelength light suppresses melatonin and decreases sleepiness with activating effects extending to sleep. This has mainly been attributed to melanopic effects, but mechanistic insights are missing. Thus, we investigated whether two light conditions only differing in the melanopic effects (123 vs. 59 lux melanopic EDI) differentially affect sleep besides melatonin. Additionally, we studied whether the light differentially modulates sensory processing during wakefulness and sleep.

Twenty-nine healthy volunteers (18-30 years, 15 women) were exposed to two metameric light conditions (high-vs. low-melanopic, ≈60 photopic lux) for 1 hour ending 50 min prior to habitual bed time. This was followed by an 8-h sleep opportunity with polysomnography. Objective sleep measurements were complemented by self-report. Salivary melatonin, subjective sleepiness, and behavioural vigilance were sampled at regular intervals. Sensory processing was evaluated during light exposure and sleep on the basis of neural responses related to violations of expectations in an oddball paradigm.

We observed suppression of melatonin by ≈14 % in the high-compared to the low-melanopic condition. However, conditions did not differentially affect sleep, sleep quality, sleepiness, or vigilance. A neural mismatch response was evident during all sleep stages, but not differentially modulated by light.

Suppression of melatonin by light targeting the melanopic system does not automatically translate to acutely altered levels of vigilance or sleepiness or to changes in sleep, sleep quality, or basic sensory processing. Given contradicting earlier findings and the retinal anatomy, this may suggest that an interaction between melanopsin and cone-rod signals needs to be considered.

**Statement of Significance:** Metameric light allows to mechanistically investigate the contribution of one specific retinal receptor. Using this approach, we here investigated the effects of high-vs. low-melanopic light for 1 hour in the evening at ecologically valid screen illuminance (≈60 photopic lux). Going beyond earlier research, we also investigated effects on sleep. We found that despite significant suppression of melatonin, other endpoints including sleep and sleep quality were not differentially affected. This underlines that melatonin suppression does not automatically translate to alterations of sleep, sleepiness, or vigilance. Further, it suggests that melanopsin effects may need to be studied in the context of cone-rod signals. Future research should thus investigate the relevance of such an interaction, which may vary between endpoints.

## Introduction

The effects of short-wavelength light on sleep and the circadian timing system are thought to be primarily mediated by specialised retinal ganglion cells that contain the photopigment melanopsin, wherefore they are intrinsically sensitive to light between 460 and 480 nm (intrinsically photosensitive retinal ganglion cells [ipRGCs]; e.g. Berson, Dunn, & Takao, 2002; Dacey et al., 2005; Lockley, Brainard, & Czeisler, 2003). Short-wavelength light exposure in the evening can acutely suppress melatonin secretion, although neuroendocrine sensitivity is characterised by considerable interindividual variability (Phillips et al., 2019; Cajochen et al., 2005; Lockley et al., 2003). Moreover, evening light exposure has been shown to improve performance on tasks requiring sustained attention acutely (e.g. Lockley et al., 2006; Cajochen et al., 2011) and increase the available processing resources on a simple cognitive task as indicated by the amplitude of the P300 component of the human event-related potential (ERP; Okamoto & Nakagawa, 2013). Further, it can decrease sleepiness in the evening, prolong sleep onset latency and consequently increase sleepiness in the morning (cf. e.g. Zeitzer, Dijk, Kronauer, Brown, & Czeisler, 2000; Chang, Aeschbach, Duffy, & Czeisler, 2015; Santhi et al., 2012; Dijk, Cajochen, & Borbély, 1991). Moreover, evaluating sleep objectively with electroencephalography (EEG), Münch et al. (2006) demonstrated that exposure to short-wavelength light compared to longer wavelengths leads to decreased slow-wave activity (SWA) during the first sleep cycle. The authors concluded this finding to reflect that the alerting effect of the light persisted into sleep, which is well in line with findings by Chellappa et al. (2013), who reported a decrease in homeostatic sleep pressure following ‘blue’ light exposure in the evening. Chang et al. (2015) additionally found the use of light-emitting e-readers to decrease and delay REM sleep, but no change in total sleep time (TST), sleep efficiency, or the duration of non-REM (NREM) sleep compared to reading a printed book.

While many of the studies point to a special relevance of the short-wavelength-sensitive melanopic system in mediating the abovementioned effects, only recently so-called metameric light conditions have allowed to specifically target specific classes of retinal receptors in selected, isolated fashion while producing no responses in other classes (Spitschan & Woelders, 2018; Tsujimura, Ukai, Ohama, Nuruki, & Yunokuchi, 2010; Estévez & Spekreijse, 1982). Such studies yield strong mechanistic insights regarding the relevance of specific receptors and thus light characteristics. Contrasting two metameric light conditions (73.5 photopic lux), that is, high- vs. low-melanopic light with 77.7 and 24.7 melanopic lux, respectively, Allen, Hazelhoff, Martial, Cajochen, and Lucas (2018) demonstrated that 5-h exposure to high-melanopic light during the evening (18:00-23:00) resulted in stronger suppression of melatonin and a decrease of subjective sleepiness in a sample of 11 volunteers. Similarly, Souman et al. (2018a) found that 3-h high-melanopic (188.8 melanopic lux, 171 lux melanopic equivalent daylight illuminance [EDI]) exposure from two hours before to one hour after habitual bed time suppressed melatonin compared with a low-melanopic (54.6 melanopic lux, 49 lux EDI; both low-/high-melanopic: 175 photopic lux) and a dim light condition, however without affecting alertness. Thus, while high-melanopic light is effective in suppressing melatonin secretion, the effects on cognition are less clear. Whether the effects of light on subsequent sleep that have been reported in other studies can largely be attributed to short-wavelength light acting on ipRGCs or rather result from an interaction with photopic or cone-mediated light characteristics is still unknown. Thus, this project aimed at investigating the effects of two metameric light conditions designed to only differ in their effects on ipRGCs but not cones on subjective and objective sleep parameters beyond melatonin suppression. We expected melatonin to be suppressed and SWA in the first sleep cycle to be decreased more effectively by high compared to low-melanopic light. Besides this, we explored the effects on sleep, sleepiness, and behavioural vigilance using self-report scales and objective measurements, respectively.

Going beyond classic, sleep parameters (i.e., SWA, sleep architecture) that describe global brain states and reflect a *macroscopic* perspective, we were also interested in whether and how light exposure alters brain processes on the *microscopic* level. More specifically, we propose this could inform whether high-melanopic artificial light exposure leads to more ‘wake-like’ processing during sleep that underlies the observed effects on ‘global brain states’ and sleep quality. To this end, we investigated whether metameric light conditions would differentially alter basic sensory processing during wakefulness and subsequent sleep. Classically, basic sensory processing has been studied using so-called oddball paradigms, where rare deviating tones are included in a sequence of frequent standard tones. These deviating stimuli are well-known to give rise to a distinct component in the human event-related potential (ERP), the so-called mismatch negativity (MMN; Sams, Alho, & Näätänen, 1983; Näätänen, Paavilainen, Tiitinen, Jiang, & Alho, 1993; for a review see Näätänen, Paavilainen, Rinne, & Alho, 2007). Although the MMN was often considered to reflect a low-level or ‘pre-attentive’ prediction error (Näätänen et al., 1993), it is nowadays best explained by an active ‘top-down’ predictive mechanism and only to a lesser extent by passive sensory adaptation and violations thereof (Wacongne, Changeux, & Dehaene, 2012). Additionally, the MMN amplitude has been shown to be modulated by (selective) attention in some studies, although not requiring (waking) attention (Woldorff et al., 1993; Alain & Woods, 1997; Woldorff, Hillyard, Gallen, Hampson, & Bloom, 1998). For instance, the MMN was found to be reduced by mental fatigue (Yang, Xiao, Liu, Wu, & Miao, 2013) and prolonged wakefulness (Raz, Deouell, & Bentin, 2001). Additionally, Vandewalle et al. (2006) reported light (during daytime) to enhance brain responses in cortical areas that support attentional oddball effects. Okamoto and Nakagawa (2013) furthermore found daytime exposure to short-wavelength light, to increase the amplitude of a later attention-modulated ERP, the P300 component. Thus, we here investigated whether 1-h pre-sleep exposure to two metameric light conditions would differentially alter attentional processes involved in early sensory processing as reflected by the mismatch response (i) during the light exposure and (ii) during a subsequent 8-h sleep opportunity (cf. Figure 1). Specifically, we hypothesised that the light exposure would differentially affect the active prediction system during sleep thus yielding stronger ERP responses in the high-melanopic condition.

**Figure 1.**
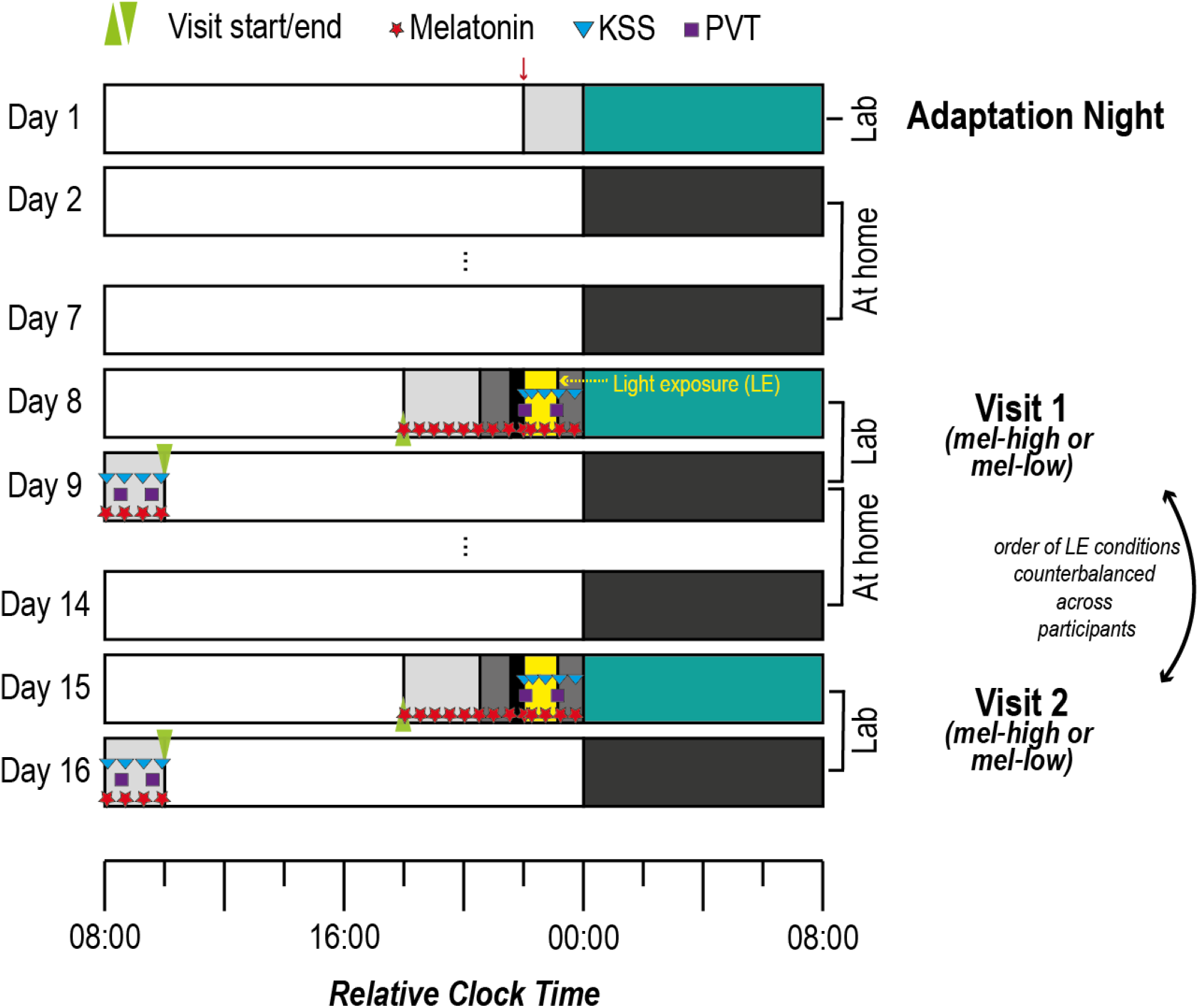
Experimental protocol for a participant with a habitual bed time (HBT) at 00:00. An adaptation night was followed by two experimental visits, which were spaced by one week and took place on the same day of the week. For 7 days before the first experimental visit and in between the two visits volunteers adhered to fixed bed and wake times to stabilise their circadian rhythms. For the experimental visits, participants arrived at the laboratory 5 h 15 min before their HBT, at the latest at 5:45 pm. From 6 pm onwards melatonin samples (red stars) were collected every 30 min. From just before the light exposure (1 h 50 min before until 50 min before HBT) until the end of the protocol, they rated their sleepiness on the Karolinska Sleepiness Scale (blue triangles) with each melatonin sample. Additionally, there were 4 psychomotor vigilance tests (PVT, 10 min). The order of the light exposure conditions (high- vs. low-melanopic) was counter-balanced across participants and sexes. The red arrow marks the point of inclusion in the study. LE = Light exposure.

## Methods and Materials

### Pre-registration

This project was pre-registered on Open Science Framework (OSF) including details about the light exposure, the analytic strategy as well as the code for the acoustic stimulation paradigm (https://www.doi.org/10.17605/OSF.IO/M23EH). As it is classified as a clinical study in Switzerland, it has also been registered on the German Clinical Trials Register (DRKS00023602). For reasons of conciseness, we will address the pre-registered analyses relating to global and omission effects as well as analyses including the data from the post-sleep recordings in (a) separate publication(s).

### Participants

Following screening procedures and after having obtained written informed consent, we had invited 40 volunteers to participate in the study between April 2019 and May 2020. Ten volunteers withdrew consent or were excluded (e.g., due to non-adherence to the study protocol) after the adaptation night or the first experimental night. The final study sample thus comprised 30 participants (15 women). Data from one male participant had to be excluded as melatonin analyses (see below) did not yield valid data (i.e., values in the *mel-high* condition already >50 pg/ml 5 h prior to habitual bed time [HBT] with no detectable rise, linear rise in the *mel-low* condition with a starting value of 16.5 pg/ml already 5 h before HBT). Thus, the final sample comprised 29 participants (15 women). German was the volunteers’ mother tongue (or comparable level) and they scored normally on questionnaires screening for sleep disorders (Pittsburgh Sleep Quality Index [PSQI], inclusion if sum values ≤ 5; Buysse, Reynolds, Monk, Berman, & Kupfer, 1989) and psychiatric symptoms (Brief Symptom Inventory [BSI]; sum scores on nine subscales and ‘global severity index’ outside clinical range; Franke & Derogatis, 2000). Furthermore, they did not habitually sleep less than 6 or more than 10 hours on work days and did not have an extreme chronotype according to the Munich Chronotype Questionnaire (MCTQ; Roenneberg, Wirz-Justice, & Merrow, 2003) or the Morningness-Eveningness Questionnaire (MEQ - German version; Horne & Östberg, 1976). The average self-selected HBT was 11:00 pm (range 10:15 pm – 00:00). Participants had to be right-handed, between 18 and 30 years of age *(M* = 23.2±2.8 years; range 19-29 years), and did not report any vision disorders nor hearing problems. They had normal or corrected-to-normal vision, and in case they wore glasses, the glasses did not have a blue light filter. All volunteers were healthy non-smokers with a body-mass-index (BMI) above 18.5 and below 25 (i.e., normal weight) and did not report any previous brain injury or relevant medical history regarding neurological or psychiatric disorders. To ensure participants did not take drugs, volunteers had to take a urine-based multi-drug panel (nal von Minden GmbH, Germany) screening for cocaine, benzodiazepines, amphetamines, tetrahydrocannabinol (THC), morphine, and methadone on the evening of every visit to the laboratory. Participants were seen by a study physician and only included if s/he did not identify reasons for exclusion. Shift-work within the three months or travelling across >2 time zones in the month prior to study participation served as additional exclusion criteria. The study protocol was approved by the ethics commission (Ethikkommission Nordwest-und Zentralschweiz; 2018-02003). Upon completion of the ambulatory and laboratory parts, participants received a remuneration of CHF 450.

### Study procedures

Data were recorded continuously between April 2019 and May 2020 with a break between mid-March and mid-May 2020 due to the Coronavirus pandemic. The experimental protocol comprised an adaptation night as well as two experimental lab visits, which were interspaced by two ambulatory phases (cf. Figure 1). All procedures were individually timed to participants’ habitual bed time. During the adaptation night, volunteers slept in the laboratory during the pre-defined habitual sleep window with an 8-h sleep opportunity with full polysomnography equipment (for details see below) to familiarise them with the laboratory setting. Sleep during the adaptation night was evaluated informally regarding sleep onset latency, sleep architecture, sleep efficiency, breathing, and limb movements to ensure that participants were indeed healthy sleepers. Sleep during this night is usually not deemed ‘representative’ and a formal evaluation comparable to the experimental nights (see below) was beyond the budget for this study. Further, melatonin was not assessed during the adaptation night as participants had not been on a circadian stabilisation protocol (see below) prior to this visit. Thus, we refrained from a formal evaluation of sleep and melatonin during the adaptation night. Following the adaptation night, volunteers kept a pre-defined constant sleep-wake cycle for the next 7 days, which was verified by sleep logs and wrist actigraphy (ActiGraph LL.C, Pensacola, FL 32502, USA). Specifically, they had to switch off lights at a specific time (±30min to the agreed [HBT]) and get up at a specific time (±30 min habitual wake time) while maintaining an 8:16 h wake-sleep cycle. In one case, the HBT was delayed by 30 minutes on the second visit as the participant had consistently gone to sleep at the end of the agreed time window for lights off. In another case, HBT was delayed due to technical failure of the screen. All participants then came to the laboratory for the first experimental visit (see below), after which they underwent the ambulatory protocol for a second time before coming to the laboratory for a second and final experimental visit.

Experimental nights took place on the same day of the week (i.e., Monday through Friday) and were usually spaced by exactly one week. In three cases, the two experimental visits were spaced by eleven, 14, and 15 days, respectively, due to unforeseen circumstances, wherefore the visits took place on a different day of the week in two cases. Only in one case, the two nights were spaced by 62 days, because the first night took place just before the laboratory was closed due to the Coronavirus pandemic and the second night took place right afterwards. Importantly, if the adaptation night and experimental visits were not exactly spaced by seven days, participants still had to adhere to a 7-day ambulatory phase to stabilise the sleep-wake cycle. We aimed at counter-balancing the order of the conditions across participants and gender subgroups. Drop-outs precluded counter-balancing among gender subgroups. Thus, the high-melanopic condition was the first for 8 men and 6 women.

#### Experimental visits

On the evenings of both experimental visits, which only differed regarding the light condition (i.e., *mel-high* [123 lux melanopic EDI] or *mel-low* [59 lux melanopic EDI], cf. Table 3), volunteers came to the lab 5 h 15 min before HBT or at 5:45 pm at the latest. Upon arrival, the room lighting was characterised by 3.2 cd/m^2^ melanopic EDL and 8.7 cd/m^2^ photopic luminance (cf. Table 1). They were served a light dinner (i.e., a sandwich) and from then on, they were seated in a comfortable chair facing the screen used for light exposure (see below), which was switched off at this time. From 6 pm (irrespective of the individual HBT), salivary melatonin samples were taken using salivettes (Sarstedt®) at 30 min intervals until the light source was switched on, 5 min into the light exposure, and again at 30 min intervals until 15 min before HBT. Saliva samples were centrifuged and frozen at −18° C immediately for later assaying. We started applying the EEG 4 h 20 min before HBT. From 3 h 20 min until 2 h 20 min prior to HBT volunteers were in dim light (0.3 cd/m^2^ melanopic EDL and 0.8 cd/m^2^ photopic luminance; see Table 1 for details and Fig. 1) listening to podcasts while keeping their eyes open. Following dim light, they were in complete darkness for 30 min with eyes open. We ensured open eyes by regularly checking the EOG for blinks and reminded participants to keep their eyes open if necessary. Ten minutes into the dark adaptation, participants performed a 10 min auditory version of the psychomotor vigilance task (PVT; Dinges, Powell, & Computers, 1985) and a 3 min resting-state EEG with open eyes, still in darkness. Note that for conciseness, we will not report the results of the resting EEG in this manuscript. Afterwards, the dim light was switched on again and served as a background light for the remainder of the evening. Just before the start of the light exposure, volunteers also completed the Karolinska Sleepiness Scale (KSS; Åkerstedt & Gillberg, 1990) for the first time. After this, they underwent 60 min of light exposure (for details on the light exposure protocol and the light conditions, please see below). During the experimental light exposure, they completed the auditory local-global oddball task (see below for details, overall duration approx. 41 minutes without breaks). This auditory task started approx. eight minutes into the light exposure following another resting-state EEG measurement to assess the acute effects of the light exposure. Further KSS ratings took place 5 minutes and again 30 minutes into the light exposure as well as just after it and 15 min before HBT. Note that the third KSS 30 minutes into the exposure was only introduced after the first 13 participants. Just after the light exposure, they completed another PVT as well as a resting-state EEG recording. During sleep, the auditory oddball task continued without interruptions. Importantly, participants were instructed to follow the instructions they had followed during wakefulness even during falling asleep and during the night. It has previously been shown that volunteers can follow such task instructions even during sleep (e.g., Andrillon, Poulsen, Hansen, Léger, & Kouider, 2016; Kouider, Andrillon, Barbosa, Goupil, & Bekinschtein, 2014). In the morning, just after having been woken up following an 8-h sleep opportunity, the volunteers resumed saliva sampling for later melatonin assays at 30 min intervals. Additionally, volunteers filled in the KSS just after waking up as well as the morning protocol of the sleep diary assessing sleep and awakening quality (Saletu, Wessely, Grünberger, & Schultes, 1987). Thirty minutes after wake-up, the morning block of assessments began. Here, two recordings of resting-state EEG activity and the completion of two more PVTs surrounded the oddball paradigm in the morning. Volunteers again indicated subjective sleepiness on the KSS after the completion of the oddball task. The morning recordings were completed under conditions of brighter light than the dim light in the evening (14.3 cd/m^2^ melanopic EDL and 38.9 cd/m^2^ photopic luminance, cf. Table 1).

**Table 1.** Overview of the radiance-derived α-opic responses (L_e_,Ω in mW/m^2^*sr) for the background light in the evening and morning and radiance-derived α-opic equivalent daylight (D65) luminances (EDL, in cd/m^2^) for the background light in the evening and morning. Photopic luminance was 8.7 cd/m^2^ (evening room), 0.8 cd/m^2^ (evening dim), and 38.9 cd/m^2^ (morning room). Radiance-derived chromaticity values (CIE 7931 xy standard observer for a 2° field) were x = 0.5 and y = 0.4 for all three background light settings. For the measurements, the measuring device was turned in the direction of the light source from the observer’s point of view. Values were calculated using the luox app (Spitschan, 2021).

|  | *S-cone-opic* |  | *M-cone-opic* |  | *L-cone-opic* |  | *Rhodopic* |  | *Melanopic* |  |
| --- | --- | --- | --- | --- | --- | --- | --- | --- | --- | --- |
| α-opic | $L_{e,\Omega}$ | EDL | $L_{e,\Omega}$ | EDL | $L_{e,\Omega}$ | EDL | $L_{e,\Omega}$ | EDL | $L_{e,\Omega}$ | EDL |
| Evening room | 1.2 | 1.5 | 9.6 | 6.6 | 14.3 | 8.8 | 5.9 | 4.0 | 4.2 | 3.2 |
| Evening dim | 0.1 | 0.2 | 0.9 | 0.6 | 1.4 | 0.8 | 0.6 | 0.4 | 0.4 | 0.3 |
| Morning room | 5.5 | 6.7 | 43.2 | 29.7 | 64.5 | 39.6 | 26.4 | 18.2 | 19.0 | 14.4 |

#### Light exposure protocol and conditions

Upon arrival at 5:45 pm or 5 h 15 min before HBT, participants were seated at a table in *evening room light* (colour rendering index [CRI CIE Ra] 94; 8.66 cd/m^2^ photopic luminance; 12 photopic lux). Starting 3 h 20 min before HBT, they were in *evening dim light* (CRI 93.88; 0.82 cd/m^2^ photopic luminance; 1 photopic lux) until HBT, except for 30 min dark adaptation from 2 h 20 min until 1 h 50 min before HBT. The dark adaptation was included to reduce potential interindividual variance <u>in the adaptation state of the eye, thereby minimising potential ‘photic history’ effects</u>. For measurements of the background light, the measuring device was turned in the direction of the light source from the observer’s point of view (i.e., 70 cm from screen at a height of 120 cm). For an overview of the radiance-derived α-opic responses and equivalent daylight (D65) luminances for the background lighting (i.e., no experimental light) please see Table 1. Suppl. Fig. 6 additionally illustrates the luminance densities for the background lighting, i.e., “evening room”, “evening dim”, and “morning room”.

From 1 h 50 min until 50 min before HBT, participants were exposed to either *high-* (CRI 7.13; 59.87 photopic lux, 122.7 lux melanopic EDI) or *low-melanopic* (CRI −112.25; 50.35 photopic lux, 59.2 lux melanopic EDI) screen light. To ensure a distance of 70 cm of the participants’ eyes from the screen, the non-moveable chair remained in a pre-specified position for the duration of the exposure and participants were asked to adopt and keep a comfortable position leaning against the backrest of the chair. Additionally, we were able to observe them using a video camera and thus correct the position if necessary. The duration of the light exposure was motivated by the aim to use a duration that would come close to for instance watching an episode of a TV series before going to bed. Additionally, a duration of one hour represented an ideal overlap with the duration of the cognitive paradigm thereby allowing to assess the acute effects of light exposure relatively close to HBT. The two conditions were matched in the activation of the L, M and S cones as quantified using CIE S026 (Commission International de l’Eclairage (CIE), 2018), which incorporate 10° Stockman-Sharpe cone fundamentals. As a consequence, the photopic illuminance, quantified using the 2° V(λ) function, was slightly different between the two conditions. Note that rods are expected to be saturated and thus not contribute to differential effects at the illuminance levels used. For a graphical illustration of the spectra, please see Figure 2 A. Please see Tables 2 and 3 for an overview of the radiance- and irradiance-derived α-opic responses and α-opic equivalent daylight (D65) irradiances/ illuminances for the two experimental screen light conditions, respectively. For the relevant photon densities, please see the supplemental material. Measures of the experimental conditions were taken at a distance of 70 cm from the screen at a height of 120 cm, that is, from the observer’s point of view. All participants underwent both experimental lighting conditions, the order was counterbalanced across participants. The light was presented on a 24-inch custom-made visual display (i.e., computer screen; Pross, 2019). This screen contains seven LEDs in total, two red LEDs with peak wavelengths (λ_p_) of 630 and 660 nm, two green LEDs (λ_p_ = 2x 520 and 550 nm) and three blue LEDs (λp = 430, 450, and 470 nm). The metameric stimuli were generated using a constrained numerical optimisation implementing the method of silent substitution (Spitschan & Woelders, 2018; Spitschan, Aguirre, & Brainard, 2015; Tsujimura et al., 2010) for non-linear light sources using Matlab’s ‘fmincon’ function (The Mathworks, Natick, MA).

**Figure 2.**
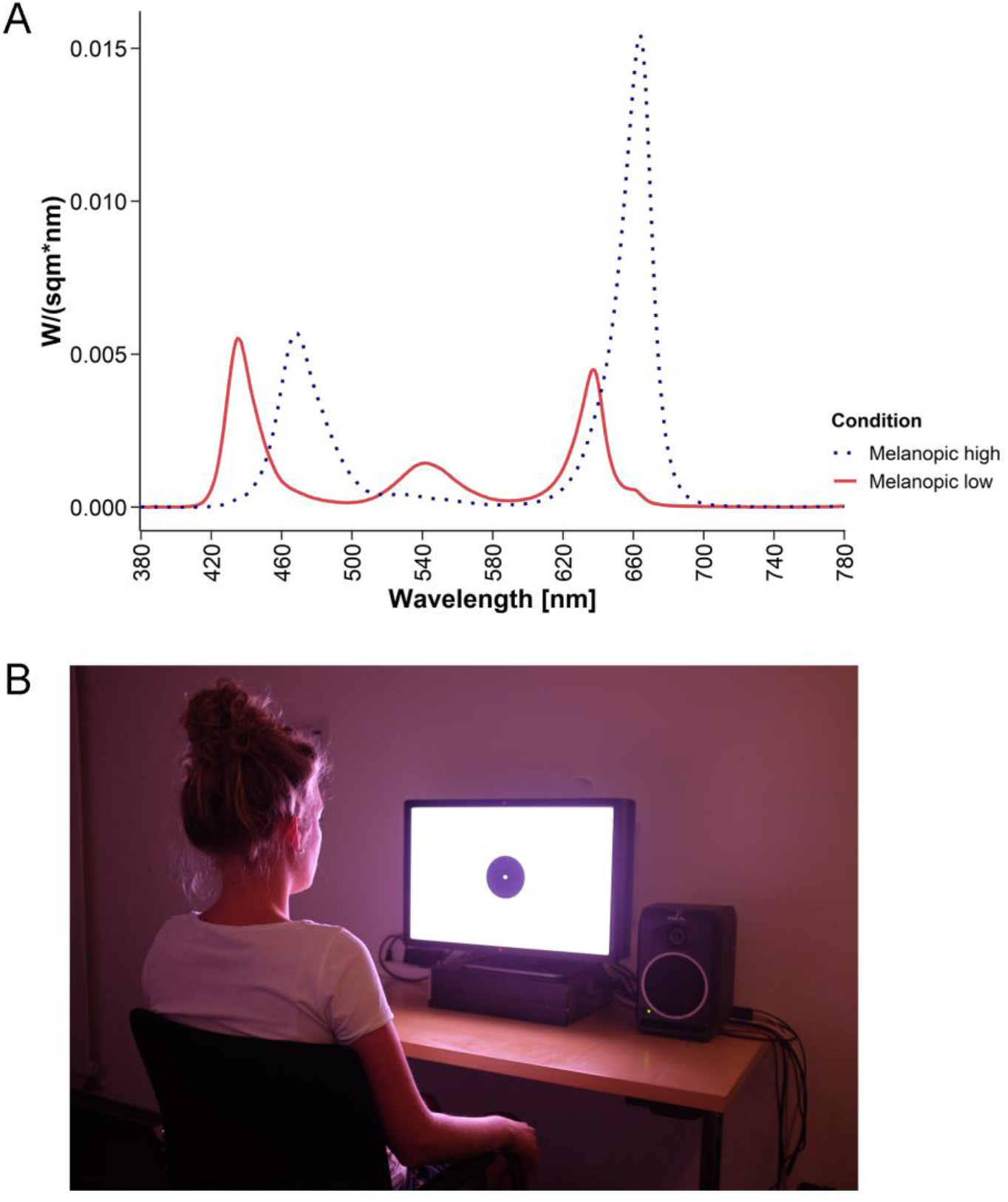
**(A)** Spectra of the two experimental screen light conditions. The mel-low condition is indicated by the solid line, the mel-high condition by the dashed line. For an overview of the spectral distributions for the two conditions reported in 1nm steps, please see the supplemental CSV file provided. **(B)** Photo illustrating the screen and the participant’s position relative to it during the light exposure. Note that the photo does not accurately reflect the colour of the screen due limitations in the reproduction of colours using uncalibrated digital RGB images. For an illustration of the irradiance-derived chromaticity coordinates (x, y) of the two light sources, please see Suppl. Fig. 1. For the (ir-) radiance-derived chromaticity values please see the captions of Tables 2 and 3.

**Table 2.** Overview of the radiance-derived α-opic responses (L_e_,Ω; in mW/m^2^*sr) for the two experimental screen light conditions and radiance-derived α-opic equivalent daylight (D65) luminances (EDL, in cd/m^2^). Photopic luminance was 10.4 cd/m^2^ (mel-low) and 8.9 cd/m^2^ (mel-high) and radiance-derived chromaticity values (CIE 1931 xy standard observer for a 2° field) were x = 0.28 and y = 0.19 for the mel-low and x = 0.29 and y = 0.17 for the mel-high condition. Measures were taken at a distance of 70 cm from the screen at a height of 120 cm, that is, from the observer’s point of view. Values were calculated using the luox app (Spitschan, 2021).

|  | *S-cone-opic* |  | *M-cone-opic* |  | *L-cone-opic* |  | *Rhodopic* |  | *Melanopic* |  |
| --- | --- | --- | --- | --- | --- | --- | --- | --- | --- | --- |
| α-opic | $L_{e,\Omega}$ | EDL | $L_{e,\Omega}$ | EDL | $L_{e,\Omega}$ | EDL | $L_{e,\Omega}$ | EDL | $L_{e,\Omega}$ | EDL |
| Mel-high screen light | 22.1 | 27.1 | 14.1 | 9.7 | 17.5 | 10.7 | 15.7 | 10.8 | 15.0 | 11.3 |
| Mel-low screen light | 20.1 | 24.6 | 14.8 | 10.1 | 17.2 | 10.5 | 27.2 | 18.7 | 31.2 | 23.5 |
| Contrast (mel-high – mel-low)/mel-low [%] | -9.3 |  | 4.7 |  | -1.8 |  | 72.9 |  | 107.3 |  |
| Ratio mel-high/ mel-low | 0.9 |  | 1.0 |  | 1.0 |  | 1.7 |  | 2.1 |  |

**Table 3.** Overview of the irradiance-derived α-opic responses (Ee; in mW/m^2^) and equivalent daylight (D65) illuminances (in lux) for the two experimental screen light conditions (background light: evening dim light). Photopic illuminance was 59.9 lux (mel-low) and 50.4 lux (mel-high) and irradiance-derived chromaticity values (CIE 1931 xy standard observer for a 2° field) were x = 0.30 and y = 0.21 for the mel-low and x = 0.30 and y = 018 for the mel-high condition. For an illustration, please see Suppl. Fig. 1. Measures were taken at a distance of 70 cm from the screen at a height of 120 cm, that is, from the observer’s point of view. Values were calculated using the luox app (Spitschan, 2021).

|  | *S-cone-opic* |  | *M-cone-opic* |  | *L-cone-opic* |  | *Rhodopic* |  | *Melanopic* |  |
| --- | --- | --- | --- | --- | --- | --- | --- | --- | --- | --- |
| α-opic | $E_e$ | EDI | $E_e$ | EDI | $E_e$ | EDI | $E_e$ | EDI | $E_e$ | EDI |
| Mel-high screen light | 103.5 | 126.7 | 79.9 | 54.9 | 96.9 | 59.5 | 142.7 | 98.4 | 162.8 | 122.7 |
| Mel-low screen light | 110.9 | 135.7 | 79.0 | 54.2 | 100.4 | 61.6 | 83.6 | 57.7 | 78.5 | 59.2 |
| Contrast (mel-high – mel-low)/mel-low [%] | -6.7 |  | 1.1 |  | -3.5 |  | 70.7 |  | 107.4 |  |
| Ratio mel-high/ mel-low | 0.9 |  | 1.0 |  | 1.0 |  | 1.7 |  | 2.1 |  |

A white background with a black spot in the centre subtending to a visual angle of 8.6° was presented on the screen to avoid a phenomenon called Maxwell’s spot, a red spot appearing in the centre of the visual field due to the absorption of light by the macular pigment (Spitschan, Jain, Brainard, & Aguirre, 2014; Gardasevic, Lucas, & Allen, 2019; Isobe & Motokawa, 1955). In the middle of the black circle, there was a small white circle, which served as a fixation point and subtended to a visual angle of 0.5°. Participants’ eyes were at a distance of 70 cm from the screen. They were instructed to lean back in the chair in a comfortable position and fixate the centre of the screen for the duration of the light exposure. We refrained from using a chin rest to increase the ecological validity of the exposure. For an illustration of the display setup, please see Figure 2B. In the morning, participants woke up and they spent the time from wake-up until the end of the protocol in *morning room light* (CRI 94.38; 38.88 cd/m^2^ photopic luminance; 55 photopic lux). A supplemental CSV file contains the spectral distributions of radiance and irradiance for the *mel-high* and the *mel-low* light conditions at each wavelength between 380 and 780 nm from the observer’s point of view, or at a distance of 20 cm from the screen for horizontal radiances. It also contains the spectral radiance distributions for the background light, that is, the *evening room light*, the *evening dim light*, and the *morning room light*. For more details see the supplemental material. Spectral measurements (range 380–780 nm; optical resolution of device 4.5 nm; reported wavelength sampling 1 nm) were performed with JETI spectroval 1501 (JETI Technische Instrumente GmbH, Jena, Germany).

#### Acoustic stimulation – Oddball paradigm

Recordings during wakefulness took place in the evening during light exposure to evaluate the acute effects of light exposure as well as in the mornings (delayed effects). We largely adopted the design that Strauss et al. (2015) had previously used, in which participants hear sequences of (in this case German) vowels (duration: 100 ms each). Note that we here focus on the early ‘local’ mismatch effects, that is, the difference between *standards* and *deviants* (cf. Figure 3). Note that acoustic stimulation paradigms of different kinds have widely been used to investigate memory consolidation and assess cognitive processing during sleep. A recent meta-analysis did not find effects on sleep architecture (Stanyer et al., 2022).

**Figure 3.**
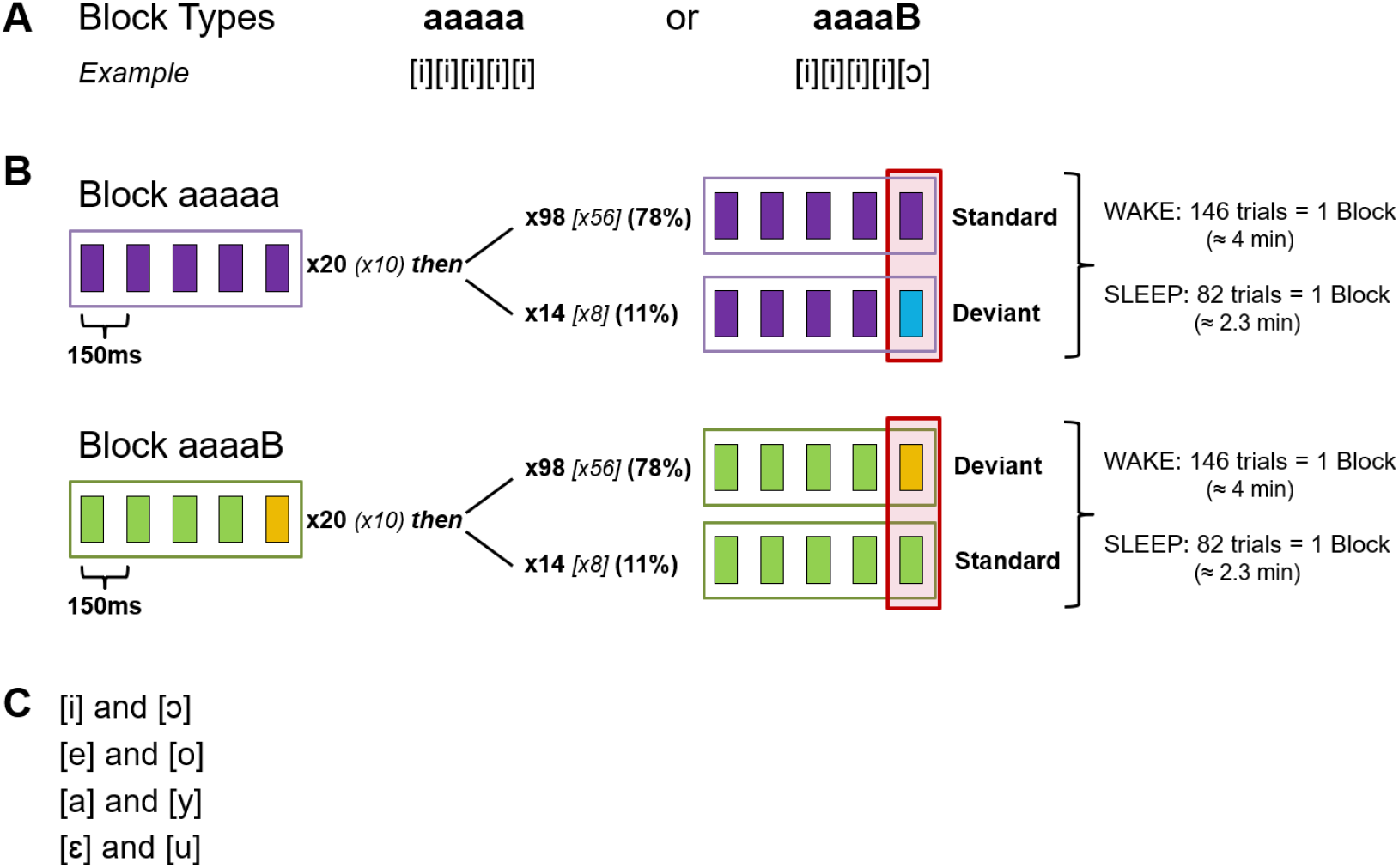
Oddball Paradigm. **(A)** Two different block types can serve as a standard. **(B)** Upper panel: In a sequence of five identical stimuli (“aaaaa ”) the fifth stimulus can be a standard (78 %) or a deviant (11 %). In “aaaaa ” blocks, participants counted the deviants. Lower panel: the standard can also be an “aaaaB” block. In this case, participants counted the standards. **(C)** Possible vowel combinations. Modified from Strauss et al. (2015). Note that the investigated effect is equivalent to the ‘local’ mismatch effect in the context of the local-global oddball. The red rectangle marks the stimuli of interest for the analyses.

In each block, one vowel was the *standard* and presented more frequently than the other, *deviant* vowel. The location of articulation of vowels that were presented in the same sequence was maximally distant (for possible combinations see Fig. 3C). Stimuli were presented in trials of five vowels, where four *standard* vowels could either be followed by another *standard* (78 % of the trials in “aaaaa” blocks, 11 % of the trials in “aaaaB” blocks) or a *deviant* (78 %in “aaaaB” and 11 % in “aaaaa” blocks, cf. Fig. 3B). Each block started with a habituation phase of 20 (10 during sleep) trials that were excluded from the analyses. In type “aaaaa” blocks, participants were instructed to count the number of *deviants* whereas in the “aaaaB” blocks they counted the number of *standards* during wakefulness and sleep. The effect of interest, the ‘local’ mismatch effect, was the comparison between deviants and standards. It was previously been shown that participants can, to some extent, also follow such instructions during sleep (Kouider et al., 2014; Andrillon et al., 2016). One block comprised 146 (±1) trials (82 [±1] during sleep). Each vowel was separated from the following one by a short gap of 50 ms and trials were separated by an interval jittered between 850-1150 ms (in 50 ms steps, average 1000 ms). Two vowel combinations were used in the evening and two during the morning recordings. During sleep, participants heard the same vowel combinations as in the evening at first, and after approx. 40 min, when we expected participants to have reached N2 sleep, the other two vowel combinations were included. Our data confirmed that participants needed an average of 10 min to reach N2 sleep (range 0.5-32.5 min). Note that besides the ‘normal’ blocks, there were also ‘control blocks’, which however are not relevant in the context of the analyses presented here. In total, participants were presented with eight standard blocks and two control blocks during wakefulness (approx. 41 min without breaks, self-paced). During sleep, there was a total of 140 standard blocks and 52 control blocks. Blocks during sleep were separated by a 6-s inter-block-interval (IBI) to mark the transition between blocks. Thus, the stimulation during sleep summed up to 7.7 h.

Before the wakefulness recordings in the evening, participants were given printed instructions explaining the structure of the paradigm and it was ensured that they understood the structure and task instructions (i.e., the stimuli they had to count). In line with previous studies (e.g., Blume, del Giudice, Wislowska, Heib, & Schabus, 2018), we individually adjusted the volume in a stepwise procedure so participants were clearly able to hear the stimuli, but at the same time could imagine falling asleep with the stimulation. A lower volume limit was pre-defined by the principal investigator according to her own experience. Additionally, as participants had to report the number of deviations after each trial, we were able to make sure that participants could indeed clearly hear the stimuli. For further details on stimulus generation and delivery, please see the supplemental material.

### Data collection and reduction

#### Electrophysiological data

We recorded the EEG signal using a BrainProducts^®^ (BrainProducts GmbH, Gilching, Germany) LiveAmp^®^ device. Data were collected at a sampling rate of 250 Hz from 23 scalp electrodes, 2 ECG, and 2 chin EMG electrodes. Eye movements were recorded from electrodes placed according to the guidelines of the American Academy of Sleep Medicine and Iber (2007) below the outer canthus of the left and above the outer canthus of the right eye. For the polysomnography setup during the adaptation night and further details please see the supplemental material. Impedances were kept below 5kΩ and checked several times throughout the recording.

For EEG analyses we used Matlab 2019a and the Fieldtrip toolbox (Oostenveld, Fries, Maris, & Schoffelen, 2010) in the version distributed by the Salzburg Brain Dynamics lab (date downloaded: 31-05-2019). For details on EEG pre-processing, the analyses of event-related potentials (ERPs) and time-frequency responses as well as analysis of EEG slow-wave activity (SWA), please see the supplemental material.

#### Sleep staging

Sleep was scored semi-automatically by the Siesta Group GmbH (Vienna, Austria; Anderer et al., 2005; Anderer et al., 2010), that is, it was scored by an algorithm and the results reviewed by a human expert. To this end, we down-sampled data from electrodes F3, F4, C3, C4, O1, and O2 as well as the EMG and EOG data acquired according to Rechtschaffen & Kales recommendations to 128 Hz.

#### Sleep cycle detection

Sleep cycles were defined according to criteria adapted from Feinberg and Floyd (1979) and as implemented in the ‘SleepCycles’ package for R (R Core Team, 2015; Blume, 2020; Blume & Cajochen, 2020). We only considered the first three sleep cycles during every night in accordance with Chellappa et al. (2013).

#### Melatonin

Salivary samples were assayed for melatonin using direct double-antibody radioimmunoassays (RIA) which has been validated by gas chromatography-mass spectroscopy with an analytical least detectable dose of 0.9 pg/ml and an analytical sensitivity of 0.2 pg/ml (NovoLytiX GmbH, Pfeffingen, Switzerland; Weber, Schwander, Unger, & Meier, 1997). Samples from two volunteers were reanalysed using an enzyme-linked immunosorbent assay (ELISA, NovoLytiX GmbH, Pfeffingen, Switzerland) as two RIA measurements had led to diverging results. In one case, the profile that better resembled the ELISA results was used for further analyses, in the other case, the average of the two measurements was used. For further details on the handling of melatonin data, please see the supplemental material.

#### Subjective sleepiness

Subjective sleepiness was repeatedly assessed with the Karolinska Sleepiness Scale (Åkerstedt & Gillberg, 1990) throughout the evenings and mornings. Specifically, participants rated their sleepiness on a 9-point Likert scale ranging from “extremely alert” to “very sleepy, great effort to keep awake, fighting sleep”.

#### Behavioural vigilance

Behavioural vigilance was assessed twice in the evenings and twice in the mornings using a 6-minute auditory version of the psychomotor vigilance task (Dinges et al., 1985; Basner & Dinges, 2011). Please note that a visual task would always have involved participants looking at a screen, which required an auditory solution. Participants had to press the space key on a keyboard in response to a 65 dB tone. The interstimulus interval varied between 2 and 9 seconds. For pre-processing, we excluded non-valid trials, that is, trials with a response time (RT) <100 ms (Basner & Dinges, 2011). We then computed the median RT as well as the fastest and slowest 10 % RTs for further processing.

#### Subjective sleep quality

Subjective sleep and awakening quality was assessed using a sleep diary by Saletu et al. (1987). More precisely, participants answered several questions in the mornings on a four-point Likert scale (“no”, “somewhat”, “moderate”, “very much”), which were then combined into the subscales “sleep quality” and “awakening quality” by averaging. Additionally, the volunteers detailed the subjectively perceived number of awakenings, and the time it took them to fall asleep.

#### Visual comfort ratings

Following the light exposure, participants also rated the screen light regarding brightness, pleasantness, glare and warmth and indicated how activating they felt the light was on a 11-step Likert scale (“not at all” to “very much”).

### Statistical analyses

All statistical analyses were done in R version 4.0.4 except for the analyses of ERPs and time-frequency analyses, which were done in Fieldtrip®. All analyses except ERP and time-frequency analyses were two-sided with the critical alpha being set to 0.05. Analyses of sleep architecture and measures of objective sleep parameters as well as visual comfort were two-sided as there were directed a priori hypotheses for a subset of analyses only relating to sleep latency. If applicable, we adjusted *p*-values for multiple comparisons using the method by Benjamini and Hochberg (1995).

#### Melatonin

For the statistical evaluation of the melatonin data, we ran three separate analyses for (i) the six measurements preceding the light exposure, (ii) the four values between the onset of the light exposure and habitual bed time (HBT), and (iii) the four assessments in the morning following wake-up. Specifically, we used advanced non-parametric analyses as implemented in the ‘nparLD’ package (Noguchi, Gel, Brunner, & Konietschke, 2012) to assess the effect of the light condition (i.e., *mel-high* vs. *mel-low*) on melatonin secretion across time. A comparison of the time-series data between conditions was followed up by comparisons at each time point. Besides time series, we also compared the area under the curve (AUC) between the conditions (for the results, please see the supplemental material). For the calculation of the AUC, we used the ‘AUC’ function implemented in the ‘DescTools’ package (Signorell & et al., 2021) with spline interpolation. We here report the ANOVA-type statistic (ATS) with degrees of freedom rounded to the next integer and relative treatment effects (RTE) as a measure of the effect size for the lighting condition effects. The RTE indicates the probability with which a randomly chosen value from the whole dataset is smaller than a randomly chosen value from a specific light condition.

#### Subjective sleepiness, behavioural vigilance, and visual comfort ratings

For the statistical analysis of subjectively reported sleepiness as assessed with the Karolinska Sleepiness Scale (KSS; Åkerstedt & Gillberg, 1990), behavioural vigilance assessed with the PVT (Dinges et al., 1985; Basner & Dinges, 2011), and visual comfort ratings for the two light sources, we also used the ‘nparLD’ package (Noguchi et al., 2012). More precisely, we evaluated condition differences and variability across time (if applicable).

For the KSS, we compared (i) the values before the light exposure, (ii) values from five minutes into the exposure until 15 minutes prior to HBT, and (iii) the values obtained in the morning. For the PVT, we compared all four assessments regarding median RT and fastest and slowest 10 % RTs. Regarding visual comfort ratings, we compared ratings obtained after light exposure to each of the two lighting conditions.

#### Sleep architecture, objective sleep parameters, subjective sleep and awakening quality

Sleep architecture, objective sleep parameters, as well as subjective sleep and awakening quality were analysed with mixed linear models as implemented in R’s ‘lme4’ package (Bates, Mächler, Bolker, & Walker, 2015). Specifically, we investigated the influence of the light exposure condition (i.e., *mel-low* vs. *mel-high;* fixed effect) and the experimental visit (i.e., *visit 1* vs. *visit 2;* fixed effect). Participant ID was modelled as a random factor. Normal distribution of the data was assessed using Shapiro-Wilk normality tests and if violated, the outcome variable was transformed to ranks using R’s ‘rank’ function prior to statistical analysis. The following variables were rank-transformed: sleep latencies, WASO, sleep efficiency, number of awakenings. For fixed effects, we report *t*-values along with degrees of freedom (*df*) rounded to the next integer as the Satterthwaite approximation results in decimals. We also report 95 % confidence interval for the fixed effects, which were computed using the ‘lme4’ bootstrapping approach. In the case of a significant interaction, this was followed with contrasts using the ‘emmeans’ package (van Lenth, 2021) with Kenward-Roger degrees of freedom and a Tukey adjustment for multiple comparisons.

#### Sleep – EEG slow-wave activity

For the comparison of SWA during NREM sleep across the first three sleep cycles, we likewise used mixed linear models based on rank-transformed data. Again, participant ID was modelled as a random factor and the two light exposure conditions as well as the cycle number (i.e., first vs. second vs. third) were included as fixed effects with an interaction between the two.

#### Electrophysiological data

The mismatch effect was investigated within each light condition and compared between the *mel-low* and *mel-high* conditions for each sleep stage. For these comparisons, we used cluster-based permutation tests as implemented in the Fieldtrip toolbox (Maris & Oostenveld, 2007).

## Results

### Melatonin

On average, melatonin values from 5 min into the light exposure until habitual bed time were lower in the *mel-high* condition than in the *mel-low* condition (*F_ATS_*(1)=6.85,*p*=.019, *RTE_mel-low_=0.537*,cf. Fig. 4A). The RTE indicates that, with a probability of 53.7 % (46.3 %), a randomly chosen value from the whole dataset was smaller than a randomly chosen value from the *mel-low* (*mel-high*) condition. Follow-up analyses indicated that melatonin values 30 minutes into (*F_ATS_*(1)=13.88,*p*<.001, *RTE_mel-low_=0.559*) as well as just after the light exposure (*F_ATS_*(1)=8.03,*p*=.012, *RTE_mel-low_=0.554*) differed significantly whereas values 5 minutes into the light exposure (*F_ATS_*(1)=0.91,*p*=0.4, *RTE_mel-low_*=0.521) and 15 min before HBT (*F_ATS_*(1)=2.93,*p*=.14, *RTE_mel-low_*=0.533) did not. Taking into account melatonin values at the two significant time points yielded an overall suppression of melatonin (i.e., difference between *mel-low* and *mel-high*) by 14.05 % (*SD* = 34.7 %). More specifically, we observed a suppression in 20 individuals, no effect of light condition in two, and, contrary to the expected effects, higher values in the *mel-high* condition, in 7 individuals (for an illustration of the individual effects see Suppl. Fig. 3). There was also a significant effect of the sampling time (*F_ATS_*(2)=31.18,*p*<.001) and a trend for a sampling time × condition interaction (*F_ATS_*(3)=2.97,*p*=.074).

Analyses yielded no condition difference among the six values preceding the light exposure (*F_ATS_*(1)=1.03,*p*=.4, *RTE_mel-low_=0.511*) nor a time × condition interaction (*F_ATS_*(5)=0.39,*p*=.8). As expected, values however varied across time (*F_ATS_*(3)=68.18,*p*<.001). Likewise, there was no difference in melatonin values assessed in the morning following wake-up (*F_ATS_*(1)=1.13,*p*=.4, *RTE_mel-low_=0.488*)or time × condition interaction (*F_ATS_*(3)=0.56,*p*=.66) but variability across time (*F_ATS_*(2)=163.01,*p*<.001). Please see Figure 4A and Suppl. Fig. 3 for a graphical illustration of the melatonin results. For additional analyses of the area under the curve (AUC), please see the supplemental material.

**Figure 4.**
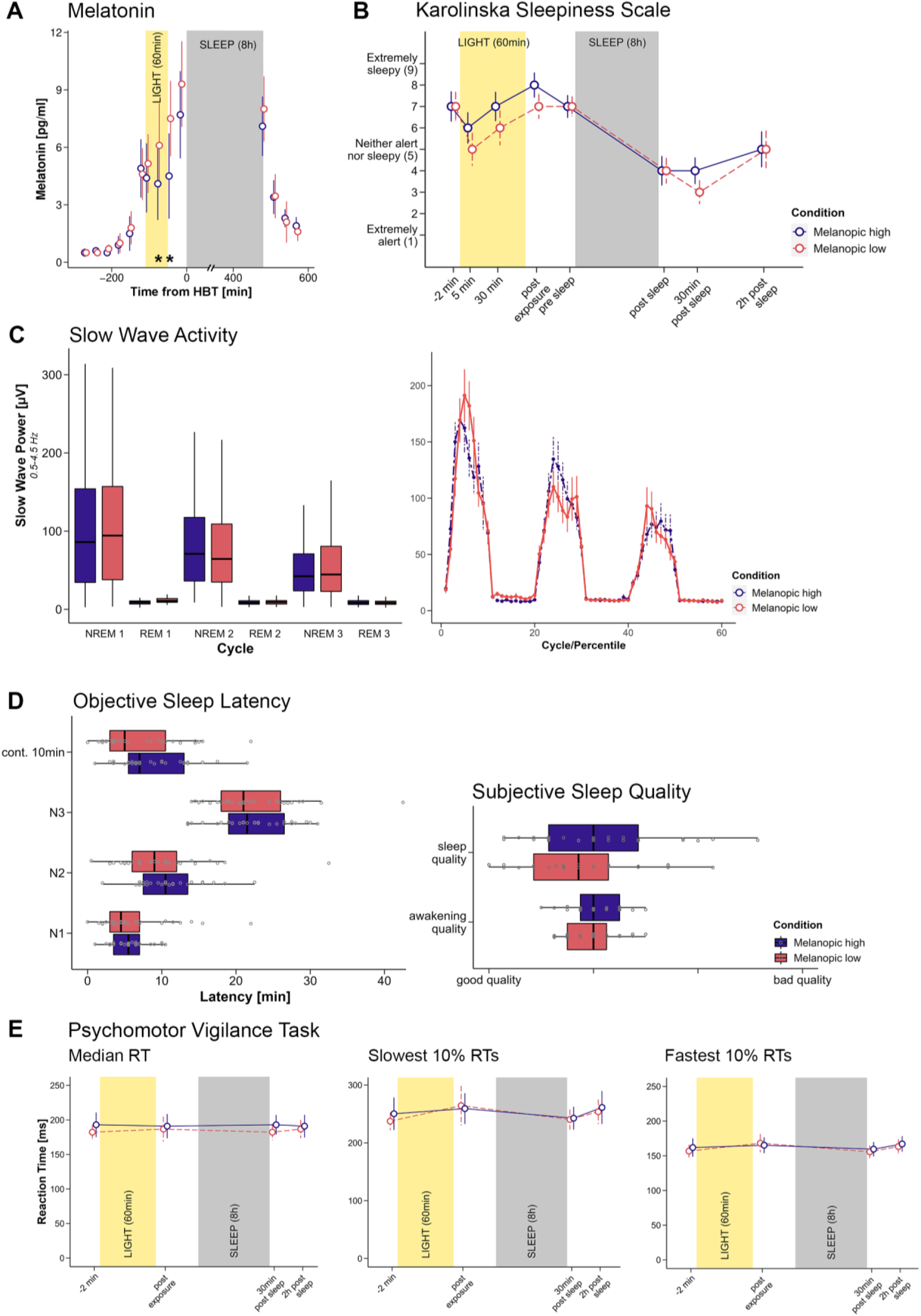
**(A)** Time course of median salivary melatonin levels (n=29, pg/ml) for the melanopic high and melanopic low condition. Vertical bars indicate the 95 % confidence intervals. The high-melanopic light was associated with a relevant suppression of melatonin secretion 30 minutes into light exposure and just after light exposure. HBT = habitual bed time. Asterisks indicate a significant difference between the light exposure conditions. **(B)** Subjective sleepiness (median and 95 % CI) assessed with the Karolinska Sleepiness Scale. There were no significant differences in sleepiness between the two lighting conditions. Please note that the third value, 30 minutes into the light exposure was only available from 17 participants as this measurement had only been introduced later. **(C)** EEG slow-wave activity (SWA) between 0.5 and 4.5 Hz across the first three sleep cycles. Left panel: Boxplots of the SWA during the NREM and REM parts of each of the first three sleep cycles. Right panel: Average power for each percentile of each NREM and REM part of a sleep cycle. Vertical bars correspond to standard errors. Generally, SWA decreased across the cycles. There were no differences between the conditions. **(D)** Objective sleep latency to continuous 10 min of sleep, N1, N2, and N3 sleep (left panel), and subjective sleep and awakening quality (right panel). The values were averaged values from subscales pertaining to sleep and awakening quality in the sleep diary by Saletu et al. (1987), respectively. There were no significant differences in self-perceived sleep or awakening quality. **(E)** Psychomotor vigilance task (PVT, 10 min auditory version). Median and 95 % CI reaction times (left panel), slowest 10 % (middle panel) and fastest 10 % (right panel) of reaction times. There were no light condition differences and no interaction between assessment point and condition. However, there was a modulation across time for all three measures (cf. main text). In boxplots, the lower and upper hinges of the boxplot correspond to the 25 % and 75 % quartiles, the thick black line indicates the median. Whiskers extend to the lowest/largest value at most 1.5× the interquartile range (IQR) from the hinges. Gray circles in boxplots represent individual values of participants. Colour code: high melanopic condition: dark purple; low melanopic condition: coral.

### Subjective sleepiness – Karolinska Sleepiness Scale

Subjective sleepiness during the light exposure and prior to HBT did not differ between the two lighting conditions (*F_ATS_*(1)=0.28,*p*=.69, *RTE_mel-low_=0.49*, cf. Fig. 4B for an illustration of the time course of KSS measurements). The RTE indicates that, with a probability of 49 % or 51 %, a randomly chosen value from the whole dataset was smaller than a randomly chosen value from the *mel-low* or *mel-high* condition, respectively. While KSS values varied across time (*F_ATS_*(2)=84.99,*p*<.001), there was no interaction between condition and time (*F_ATS_*(2)=1.36,*p*=.36). For the individual differences between the two light exposure conditions five and 30 minutes into as well as just after the light exposure, please see Suppl. Fig. 4. As expected, KSS ratings before the beginning of the light exposure did not differ between the two lighting conditions (*F_ATS_*(1)=0.12,*p*=.73, *RTE_mel-low_=0.488*). Likewise, there was no condition difference in the morning (*F_ATS_*(1)=2.11,*p*=.34, *RTE_mel-low_=0.477*). While KSS values varied across time in the morning (*F_ATS_*(1)=14.89,*p*<.001), there was no condition × time interaction (*F_ATS_*(2)=1.66,*p*=.34) either.

### Behavioural Vigilance – Psychomotor Vigilance Task

There were no condition differences regarding the median reaction time across the four measurement points (*F_ATS_*(1)=0.73,*p*=.44, *RTE_mel-low_=0.492*) with RTE suggesting that, with a probability of 49.2 % (50.8 %), a randomly chosen value from the whole dataset was smaller than a randomly chosen value from the *mel-low* (*mel-high*) condition. While RTs varied across time (*F_ATS_*(1)=12.14,*p*<.001), there was no time × condition interaction (*F_ATS_(2)=1.03, p*=.44, cf. Fig. 4E). Specifically, RT slowed from before (*mdn* = 182.25 ms) to after light exposure (*mdn* = 193 ms; *F_ATS_*(1)=14.62, *p*<.001), became shorter in the morning again (*mdn* = 186.75 ms; *F_ATS_*(1)=15.7, *p*<.001) and slowed again until after the completion of the local-global oddball paradigm (*mdn*= 191.00 ms; *F_ATS_*(1)=17.47, *p*<.001).

Likewise, there was no condition difference for the 10 % fastest RTs (*F_ATS_*(1)=1.09,*p*=.38, *RTE_mel-low_=0.491*) and no time × condition interaction (*F_ATS_*(3)=1.72,*p*=.22), but RTs varied across time (*F_ATS_*(2)=9.31,*p*<.001, cf. Fig. 4E). RTs among the fastest 10 % of the trials were again faster before the light exposure (*mdn* = 151.75 ms) than afterwards (*mdn* = 160.3 ms; *F_ATS_*(1)=13.31, *p*<.001). In the morning, volunteers’ reaction times were again faster than in the evening (*mdn* = 155.65 ms; *F_ATS_*(1)=29.54, *p*<.001), and they slowed again in the morning from before to after the oddball paradigm recording (*mdn* = 158.0 ms; *F_ATS_*(1)=11.54, *p*=.001).

In line with these results, also the 10 % slowest RTs did not differ between lighting conditions (*F_ATS_*(1)=0.09,*p*=.8, *RTE_mel-low_=0.497*) and there was no time × condition interaction (*F_ATS_*(2)=0.28,*p*=.8). Again, the results confirmed a variability of the 10 % slowest response times throughout the experimental visit (*F_ATS_*(2)=11.79,*p*<.001, cf. Fig. 4E). Specifically, we observed a slowing of RTs throughout the evening light exposure (*mdn_pre_*= 227.25 ms; *mdn_post_*= 242.60 ms; *F_ATS_*(1)=8.59, *p*=.005), RTs became again faster following sleep (*mdn* = 230.39 ms; *F_ATS_*(1)=21.76, *p*<.001) and then again slowed until the end of the experimental visit in the morning (*mdnpost* = 243.9 ms; *F_ATS_*(1)=30.65, *p*<.001).

### Visual comfort

Participants perceived the *mel-high* condition as brighter than the *mel-low* condition (*F_ATS_*(1)=4.62,*p*=.031, *RTE_mel-low_=0.432*, cf. Suppl. Fig. 2 for an illustration of the subjective ratings). There were no conditions differences in the perceived warmth of the light (*F_ATS_*(1)=0.34,*p*=.56, *RTE_mel-low_*=0.517), its pleasantness (*F_ATS_*(1)=0.1,*p*=.75, *RTE_mel-low_=0.51*), its glare (*F_ATS_*(1)=2.53,*p*=.11, *RTE_mel-low_*=0.451), or regarding how activating it was perceived (*F_ATS_*(1)=0.04,*p*=.85, *RTE_mel-low_*0.507).

### Subjective sleep and awakening quality

Subjective sleep quality as assessed on a Likert-Scale from 1-4, where 1 denotes “good quality” and 4 denotes “bad quality did not differ between conditions (*b* = 0.11, *t*(46) = 0.48, *p* = .64; *mdn_mel-low_* = 0.86, *mdn_mel-high_ = 1.0*) nor the visit (*b* = −0.06, *t*(46) = −0.26,*p* = .79; *mdn_v1_* = 1.0, *mdn_v2_* = 1.0), neither was there an interaction between condition and visit (*b* = 0.14, *t*(27) = 0.37, *p* = .71). Likewise, awakening quality did not differ between light exposure conditions (*b* = 0.5, *t*(34) = 1.69,*p* = .1; *mdn_mel-low_* = 1.0, *mdn_mel-high_* = 1.0), nor between visits (*b* = 0.12, *t*(34) = 1.49, *p* = .15; *mdn_v1_* = 1.0, *mdn_v2_* = 1.0), nor was there an interaction between condition and visit (*b* = −0.21, *t*(27) = −1.33, *p* = .19). Supplementary Tables S1 and S2 provide a detailed overview of the results and Fig. 4D (right panel) a graphical illustration.

### Objective sleep parameters & sleep architecture

All participants were healthy sleepers, which was ensured during an adaptation night (see above). Differences in latency to continuous 10 minutes of sleep did not differ between light exposure conditions (*b* = −9.52, *t*(50) = −1.61, *p* = .12; *mdn_mel-low_* = 5 min, *mdn_mel-high_* = 7 min; cf. Fig. 4D), but latency was decreased at the second experimental visit (*b* = −12.96, *t*(50) = −2.2, *p* = .033; *mdn_V1_* = 10 min, *mdn_V2_* = 5.5 min). There was no condition × visit interaction (*b* = 3.8, *t*(27) = 0.4, *p* = .69). Latency to N2 tended to be shorter in the *_mel-low_* condition (N2: *b* = −11.27, *t*(48) = −1.93, *p* = .06; *mdn_mel-low_* = 9 min, *mdnmel-high* = 10.5 min), and at the second visit (N2: *b* = −17.03, *t*(48) = −2.92, *p* = .006; *mdn_V1_* 10.5 min, *mdn_V2_* = 7.5 min) but there was no interaction (N2: *b* = 10.7, *t*(27) = 1.11, *p* = .28). Latency to N1, N3, or REM did not differ between conditions (N1: *b* = −2.75, *t*(46) = −0.43, *p* = .66; N3: *b* = 2.05, *t*(45) = 0.33, *p* = .74; REM: *b* = 1.08, *t*(53) = 0.17, *p* = .86; cf. Fig. 4D) or experimental visits (N1: *b* = −4.15, *t*(46) = −0.66, *p* = .52; N3: *b* = −6.69, *t*(45) = −1.09, *p* = .28; REM: *b* = 5.98, *t*(53) = 0.94, *p* = .34) nor was there an interaction (N1: *b* = −1.11, *t*(27) = −0.1, *p* = .92; N3: *b* = −6.08, *t*(27) = −0.59, *p* = .56; REM: *b* = −9.6, *t*(27) = −1, *p* = .32). Table 4 and supplementary table S3 provides an overview of the results.

**Table 4.**
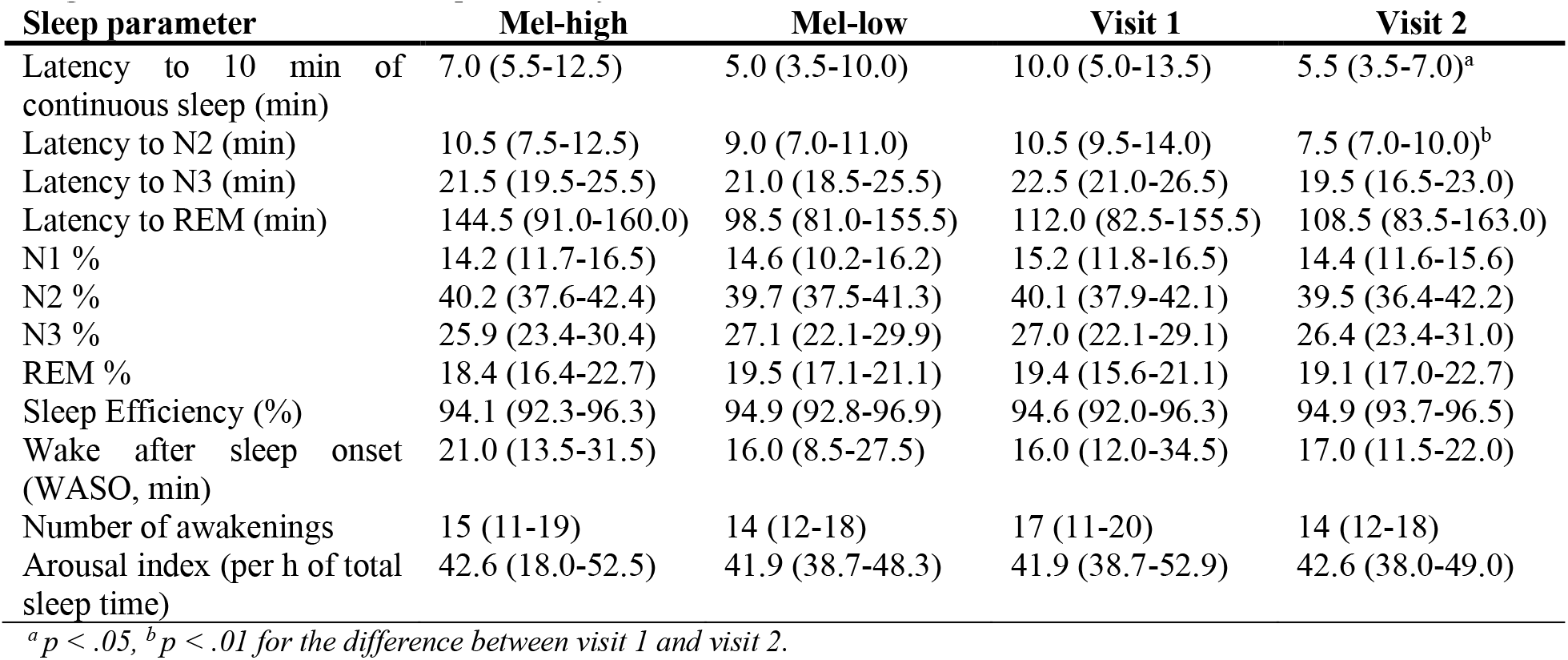
Overview of the median and 95% confidence intervals of for latency to 10 min of continuous sleep, latency to N2, N3, and REM, percentage of N1, N2, N3, REM sleep, and wakefulness, sleep efficiency, wake after sleep onset (WASO), the total number of awakenings, and the arousal index (i.e., number of arousals per hour of total sleep time) for each condition and visit. Calculations have been performed using R’s ‘DescTools ’ package (Signorell & et al., 2021), the reported confidence intervals are two-sided.

Percentages of N1, N2, N3, or REM did not differ between conditions (N1: *b* = −1.85, *t*(33) = - 0.93, *p* = .36; N2: *b* = 0.59, *t*(40) = 0.28, *p* = .78; N3: *b* = 2.65, *t*(34) = 1.14, *p* = .26; REM: *b* = −1.38, *t*(40) = −0.76, *p* = .46) or experimental visits (N1: *b* = −2.17, *t*(33) = −1.09, *p* = .28; N2: *b* = 0.40, *t*(40) = 0.19, *p* = .84; N3: *b* = 2.26, *t*(34) = 0.97, *p* = .34; REM: *b* = −0.49, *t*(40) = −0.27, *p* = .78) nor was there an interaction (N1: *b* = 2.17, *t*(27) = 0.57, *p* = .86; N2: *b* = −1.12, *t*(27) = −0.3, *p* = .76; N3: *b* = −3.83, *t*(27) = −0.88, *p* = .38; REM: *b* = 2.77, *t*(27) = 0.86, *p* = .40). An overview of the results can be found in table 4 and suppl. table S4.

Wake after sleep onset (WASO) did not differ between conditions (*b* = 5.96, *t*(39) = 0.98, *p* = .34) or visits (*b* = 6.13, *t*(39) = 1.0, *p* = .32) following rank transformation. There was no condition × visit interaction (*b* = −18.99, *t*(39) = −1.73, *p* = .096). For an overview of the results for WASO see table 4 and suppl. table S5.

Neither did sleep efficiency differ between conditions (*b* = −7.44, *t*(39) = −1.23, *p* = .22) or visits (*b* = −5.34, *t*(39) = −0.92, *p* = .36). There was no condition × visit interaction (*b* = 20.51, *t*(27) = 1.88, *p* = .072). Table 4 and suppl. table S6 provides an overview of the results for sleep efficiency.

Also the number of awakenings did not differ between conditions (*b* = 0.61, *t*(36) = 0.1, *p* = .92) or visits (*b* = −1.45, *t*(36) = −0.23, *p* = .82) and there was no interaction (*b* = −3.73, *t*(27) = −0.32, *p* = .76). Likewise, for the arousal index, that is, the number of arousals per hour of total sleep time, there was no difference between conditions (*b* = −0.63, *t*(31) = −0.15, *p* = .88) or visits (*b* = −1.42, *t*(31) = −0.34, *p* = .74) and there was no condition × visit interaction (*b* = 0.29, *t*(27) = 0.036, *p* = .98). For an overview of the results for the number of awakenings and the arousal index see table 4 and supplementary tables S7 and S8, respectively.

### EEG slow-wave activity

There was no difference in SWA (0.5-4.5 Hz) between the two light exposure conditions (*b* = 52.45, *t*(1708) = 0.75, *p* = .46), neither was there an interaction between condition and cycle number (i.e., first, second, or third; *b* = −28.37, *t*(1708) = −0.87, *p* = .38). As expected, SWA decreased across the first three cycles (*b* = −165.4, *t*(1708) = −7.2,*p* < .001). For an overview of the results for SWA see Fig. 4C and supplementary table S9.

### Event-related potentials (ERPs)

Please note that additional time-frequency analyses confirmed the ERP results detailed below. For the results of these analyses, please see the supplemental material.

#### Wakefulness

During wakefulness, there was a significant mismatch effect with deviants resulting in stronger early (mel-low: 52-148 ms, *p* = .004; mel-high: 52-148 ms, *p* = .004) as well as late (mel-low: 160-424 ms, *p* < .001; mel-high: 160-444 ms, *p* < .001) responses (cf. Fig. 5 A/F). There were no differences between the two light exposure conditions (all *p* > .28).

**Figure 5.**
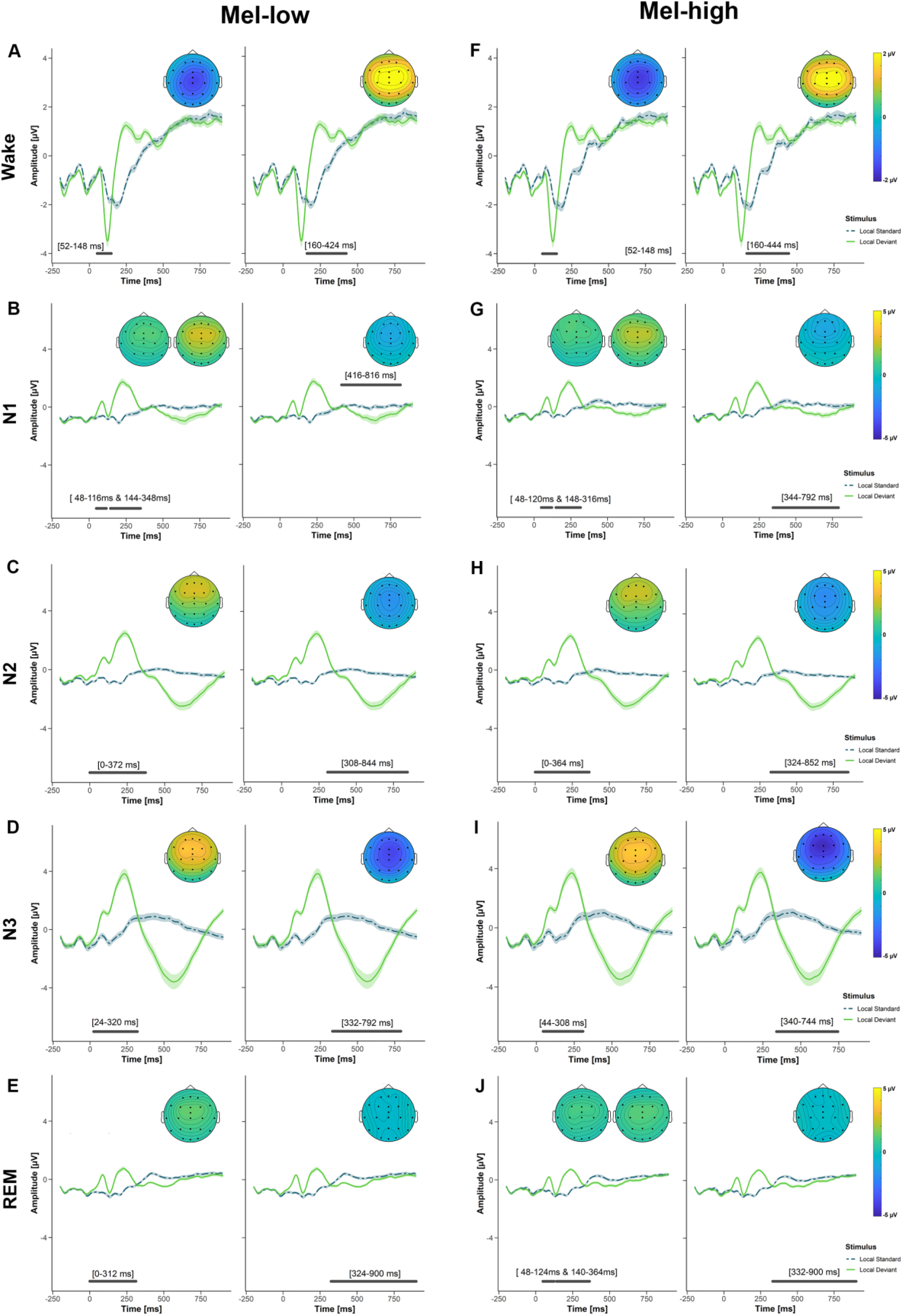
Event-related potential (ERP) effects for the mismatch effect during wakefulness (A, F), N1 (B, G), N2 (C, H), N3 (D, I), and REM (E, J) in the mel-high and mel-low conditions. The plots show the average potential across all electrodes that were part of the significant cluster. The shaded area corresponds to the standard error. Significant electrodes are indicated by black dots in the topoplots. The topoplots show the difference between the ERPs evoked by deviants and standards for each significant cluster. Significant time windows are indicated by the grey horizontal bars included in each figure. Only significant clusters are shown (cf. main text for more information).

#### Sleep

##### N1

During N1, there was a mismatch effect with an early (mel-low: 48-116 ms, *p* = .029 and 144-348 ms, *p* < .001; mel-high: 48-120 ms, *p* = .029 and 148-316 ms, *p* < .001) and a late (mel-low: 416-816 ms, *p* < .001; mel-high: 344-792 ms, *p* < .001) component (cf. Fig. 5B/G). There was no difference between the two light conditions (all *p* > .22).

##### N2

During N2, the mismatch effect persisted with early (mel-low: 0-372 ms, *p* < .001; mel-high: 0-364 ms, *p* < .001) and late components (mel-low: 308-844 ms, *p* < .001; mel-high: 324-863 ms, *p* < .001; cf. Fig. 5 C/H). There were no differences between the two light exposure conditions (all *p* > .5).

##### N3

The mismatch effect continued to be present with an early (mel-low: 24-320 ms, *p* = .035; mel-high: 44-308 ms, *p* < .001) and a late cluster (mel-low: 332-792 ms, *p* < .001; mel-high: 340-744 ms, *p* < .001; cf. Fig. 5 D/I). There was no difference between the two light conditions (no clusters).

##### REM

For the mismatch effect, there was again an early (mel-low: 0-312 ms, *p* < .001; mel-high: 48-124 ms, *p* < .001 and 140-364 ms, *p* < .001) and a late (mel-low: 324-900 ms, *p* < .001; mel-high: 332-900 ms, *p* < .001; cf. Fig. 5 E/J) component during REM. There was a difference between the light exposure conditions (*p* = .038) with the mel-low condition being associated with a larger amplitude between 0 and 92 ms above frontocentral electrodes.

## Discussion

One-hour pre-sleep exposure to two metameric lighting conditions that specifically targeted intrinsically photosensitive retinal ganglion cells (ipRGCs; melanopic ratio 2.1, photopic illuminance ≈ 60 lux) differentially affected melatonin secretion. As expected, high-melanopic light acutely suppressed melatonin more effectively than low-melanopic light resulting in an average relative suppression of about 14 %. Going beyond earlier studies, using metameric light, we also assessed the effects of pre-sleep light exposure on self-reported sleep quality as well as objective PSG-derived sleep variables. Sleep did not differ between the two light conditions despite their neuroendocrine effectiveness. In line with this, the two light conditions did not differentially modulate subjective sleepiness or behavioural vigilance. We also found that basic sensory processing of a neural mismatch response was retained during all sleep stages but likewise not differentially affected by the light conditions. Our findings underline that melatonin suppression does not automatically translate into alterations of sleepiness or sleep, as well as the notion that other light characteristics besides melanopic effects may be involved in modulating the effects of evening light beyond the neuroendocrine response (Revell, Arendt, Fogg, & Skene, 2006; Brown, Thapan, Arendt, Revell, & Skene, 2021).

At first sight, melatonin suppression by about 14% seems rather little. The high-melanopic light was more effective at suppressing melatonin than the low-melanopic light in 20 out of 29 participants (average suppression 30.2%) with no change in two and reversed pattern in seven participants (for more details see supplemental material). Given the considerable differences in sensitivity to light exposure, such a result seems expectable (Phillips et al., 2019). Furthermore, studies and thus also the magnitude of effects are often difficult to compare directly due to differences in for instance the duration and/or the timing of light exposure, pupil state (i.e., dilated vs. non-dilated), prior light history, and light characteristics (i.e., photopic illuminance, melanopic effectiveness). Additionally, the choice of the condition which experimental melatonin values are compared to is crucial. Especially when light exposure is compared to dim light, the resulting suppression rates are very high (e.g., Zeitzer et al., 2000; Thapan, Arendt, & Skene, 2001). Here, we aimed at comparing two ecologically valid light exposure scenarios, that is, exposure for 1 hour at usual screen illuminance (approx. 60 photopic lux) with a melanopic ratio of 2.1, which likely decrease the effect size.

Although it is similarly difficult to compare other acute or delayed (i.e., subsequent to light exposure) effects besides melatonin suppression to results obtained in other studies for the abovementioned reasons, their absence in the present study is somewhat astonishing. With some exceptions (e.g., Higuchi, Motohashi, Liu, & Maeda, 2005), pre-sleep light exposure has relatively consistently been associated with reduced subjective sleepiness and increased behavioural vigilance, longer sleep onset latencies, and the suppression of melatonin secretion (Chang et al., 2015; Cajochen, Kräuchi, Danilenko, & Wirz-Justice, 1998; for a review see Souman, Tinga, te Pas, van Ee, & Vlaskamp, 2018b; Gooley et al., 2011; Chellappa et al., 2011). In particular, the suppression of melatonin and possible downstream effects such as delays in sleep onset (Santhi et al., 2012; 4 h light exposure until 25 min before HBT) or alterations in sleep architecture and homeostatic sleep pressure (Münch et al., 2006; light exposure for 2 h ending 1.25 h after HBT), have largely been attributed to the melanopic rather than the cone-mediated system. This notion has received support from findings that melatonin suppression is particularly strong when short wavelength proportions are high (Cajochen et al., 2005; Chellappa et al., 2011; Revell et al., 2006) and that ‘blue’-blocking glasses can mitigate neuroendocrine and alerting responses (van der Lely et al., 2015). A recent meta-analysis even confirmed that the relationship between melanopic illuminance and the delay in sleep onset follows a dose-response relationship (Giménez et al., 2022). Further mechanistic evidence comes from studies using metameric conditions designed to affect the melanopic system only, as in the present study (Allen et al., 2018; Souman et al., 2018a). However, comparing conditions with high vs. low power between 450-500 nm at constant photopic illuminance (175 lux), Souman et al. (2018a) likewise found no differential effects on vigilance levels despite melatonin suppression reaching almost 50 % (55 vs. 189 melanopic lux and 49 vs. 171 lux melanopic EDI [melanopic ratio 3.5] in the *mel-low* and the *mel-high* conditions, respectively). A rather weak relationship between melatonin suppression, alertness, and performance has also been reported in other studies (Rüger, Gordijn, Beersma, De Vries, & Daan, 2005; Kayumov et al., 2005) and Lok, van Koningsveld, Gordijn, Beersma, and Hut (2019) showed that light and melatonin affected behavioural vigilance independently, at least during daytime. In their study, melatonin ingestion in the afternoon increased sleepiness, but bright light exposure did not alter sleepiness nor behavioural vigilance. Future studies will have to verify whether this independence also holds true for the relationship between endogenous melatonin and light exposure in the evening. Taking this notion even a step further, whether vigilance, subjective sleepiness, melatonin secretion and eventually sleep may be sensitive to different light characteristics, is still an open question. Another reason for the absence of effects on subjective sleepiness ratings or vigilance as assessed with the PVT could be that the contrast in melanopic EDI was not strong enough or may require longer light exposure durations. In particular, in the study using metameric light conditions by Allen et al. (2018), participants underwent 5 hours of light exposure while the contrast between low- and high-melanopic conditions was 24.7 vs. 77.7 melanopic lux, and light’s alerting and melatonin supressing effects only appeared in the last third of the light exposure. Likewise, also the effects on slow-wave activity may be sensitive to exposure duration (Münch et al., 2006; Chellappa et al., 2013: both 2 hours, conditions were not metameric though), which we here found to be unmodulated by prior light exposure. In the study by Münch et al. (2006), light exposure also lasted until 1.25 h beyond habitual bedtime. In sum, one-hour exposure prior to HBT as in the present study may have been too short for differential effects beyond acute melatonin suppression to occur. Especially regarding sleep, acute effects may also not have been long-lived enough. At least, differences in melatonin were levelled again 15 min before lights off, that is, 35 min after the end of the light exposure.

Besides this, cone and rod signals may well be relevant in addition to melanopic effects. As ipRGCs receive cone and rod inputs, it is plausible that the effects seen here are due to an interaction with melanopsin signals. In line with this, a recent study reported that behavioural vigilance (i.e., hit rate) was improved by both blue and red light compared to dimmer white light in day- and night shift workers (Figueiro & Pedler, 2020). The exact nature of this potential interaction between different light characteristics remains to be clarified in future studies. Whether and how metameric light affects sleep beyond melatonin suppression has, to the best of our knowledge, not previously been investigated. While it has been known for a long time that melatonin is neither sufficient nor necessary for sleep (Cajochen, Kräuchi, & Wirz-Justice, 2003; Rüger et al., 2005), our finding again underlines that sleep and melatonin suppression are not necessarily linked.

Regarding basic sensory processing as reflected by the mismatch response, we did not find evidence for differential effects of the low- vs. high-melanopic light exposure conditions. This is despite earlier research suggesting that particularly the melanopic photoreceptor system may mediate effects of short-wavelength light on cognition (Vandewalle et al., 2007). However, considering the absence of differential effects on sleepiness, vigilance, or sleep apart from melatonin suppression, the absence of such effects may not be particularly surprising. Moreover, although bright light enhances responses in areas supporting attentional oddball effects (Vandewalle et al., 2006) and can affect later components such as the P300 (Okamoto & Nakagawa, 2013), especially early responses such as a mismatch effect may be relatively insensitive to rather subtle variations in the available processing resources resulting from light exposure. Future studies should evaluate whether later processing stages are sensitive to the effects of light and whether larger contrasts or variations of the effects on other retinal receptors are involved. Besides the absence of lighting-related differential effects, we found the mismatch response to be retained during all sleep stages, although the shape of the ERP considerably changed when participants fell asleep. This is in contrast to previous research by Strauss et al. (2015), who used the same auditory oddball paradigm during an afternoon nap, and who had concluded that sleep disrupts the mismatch response leaving only (passive) sensory adaptation mechanisms intact. Generally, with a high number of trials across a whole night and thus an excellent signal-to-noise ratio (SNR), we observed a stronger K-complex (KC)-like response elicited by deviants compared to standards, most prominently during N2 and deep N3 sleep. During N1 and REM, the effect was less strong albeit still present, which may be due to a decreased SNR in these sleep stages due to the presence of eye movements and larger trial-to-trial variability of the EEG signal. Interestingly, the observed pattern in ERPs as well as time-frequency analyses is very similar to what has been reported to be evoked by salient or subjectively relevant stimuli such as one’s own name or an unfamiliar voice (Blume et al., 2016; Blume et al., 2018). This suggests that deviants continued to be salient during all sleep stages.

Several limitations have to be considered that at the same time stake out the scope for future research. First, the sample only comprised young healthy sleepers. With increasing age or the presence of vulnerability factors, the sensitivity to light may well change. Thus, the results presented here should not be generalised beyond the investigated sample. Moreover, as participants only reported to the lab in the early evening, we cannot exclude that the light history during the day had an effect. This could be relevant as it has previously been shown that bright light during the day may decrease the susceptibility to light in the evening (Rångtell et al., 2016; Smith, Schoen, & Czeisler, 2004; Hébert, Martin, Lee, & Eastman, 2002). In our sample though, the laboratory conditions took place seven days apart in most participants, wherefore seasonal differences are unlikely. Furthermore, the time they reported having spent under the open sky before coming to the lab did not differ between the two conditions (*mel-high*: 102.4 min ± 101.9; *mel-low:* 99.1 min ± 72.6; *t*(28) = 0.62,*p* = .8). Likewise, there was no difference in the perceived brightness on a scale from 1 (very cloudy day) to 10 (bright summer’s day) during this time (*mel-high:* 5.5 ± 2.7; *mel-low:* 5.8 min ± 2.6; *t*(28) = −0.41, *p* = .68). Thus, it seems unlikely that the individual prior light history affected our results.

To conclude, using two metameric light conditions that exclusively differ in their effects on ipRGCs by a factor of 2.1x, we find that early sensory processing was not differentially modulated, neither during wakefulness nor sleep. Beyond this, our findings support the notion that differences in the acute suppression of melatonin in strictly controlled light settings do not automatically translate to differences in altered levels of behavioural vigilance or experienced sleepiness and neither to differential changes in sleep or sleep quality. Thus, neuroendocrine and sleep-related mechanisms are not proxies of each other and should be investigated as such. Last, this suggests that an interaction between melanopsin and cone-rod signals may be involved in the occurrence of such effects.

## Ethical approval

Ethical approval was provided by the cantonal ethics commission (Ethikommission Nordwest-und Zentralschweiz; 2018-02003). The study was conducted in accordance with Swiss law and the Declaration of Helsinki.

## Author Contributions

CB conceptualised the project, supported by MN, TB, MS, and CC. CB acquired the data, conducted the analyses, and wrote the first draft of the manuscript. All authors provided critical review of the manuscript and approved the submitted version.

## Acknowledgements

We thank all participants, who volunteered to participate in the project. We especially also thank the interns and master students Anaité del Río, Patricia Egli, Corinna M. Hofer, Jessica Jacobs, Melina Koller, Daniela Lindegger, Marlene Schmidt, and Natascha Stoffel without whom the study could not have been conducted. We also thank our colleague Isabel Schöllhorn for her valuable expertise and help with measuring the light conditions, Oliver Stefani for the help with the computation of the photon densities, and Corrado Garbazza for his help with the medical screening of participants.

