## Supplemental Material for "Melatonin suppression does not automatically alter sleepiness, vigilance, sensory processing, or sleep"


Christine Blume[1,2] [0000-0003-2328-9612], Maria Niedernhuber[3] [0000-0002-4459-6796],
Manuel Spitschan[1,2,4] [0000-0002-8572-9268], Helen C. Slawik[5], Martin P. Meyer[5] [0000-0002-8041-5884],
Tristan A. Bekinschtein[3]§ [0000-0001-5501-8628], Christian Cajochen[1,2]§ [0000-0003-2699-7171]

§ These authors contributed equally

[1]Centre for Chronobiology, Psychiatric Hospital of the University of Basel, Basel, Switzerland (institution, where the work was performed)

[2]Transfaculty Research Platform Molecular and Cognitive Neurosciences, University of Basel, Basel, Switzerland

[3]Consciousness and Cognition Lab, Department of Psychology, University of Cambridge, Cambridge, United Kingdom

[4]Translational Sensory and Circadian Neuroscience, Max Planck Institute for Biological Cybernetics, Tübingen, Germany

[5]TUM Department of Sport and Health Sciences (TUM SG), Technical University of Munich, Munich, Germany

[6]Psychiatric Hospital of the University of Basel, Basel, Switzerland

**Corresponding Author:**

Dr. Christine Blume

Centre for Chronobiology

Psychiatric Hospital of the University of Basel

Wilhelm-Klein-Str. 27

CH-4002 Basel, Switzerland

### Supplemental Methods

*Stimulus generation*

Vowels for the auditory stimuli were uttered by a female human native German speaker, recorded with a Zoom H2n recording device, and pre-processed in Audacity 2.3.0®. More specifically, following denoising each vowel was cut to 100 ms duration with a 10 ms fade-in and 10 ms fade-out phase. We then applied a low-pass filter at 5000 Hz (6 dB) and normalised the stimuli to a peak amplitude from -1 to 0 individually for each stereo channel and removed direct current (DC). Vowels were concatenated to yield the sequences of four or five stimuli, where each stimulus was 100 ms long with a 50 ms inter-stimulus interval (i.e., silence). Afterwards, the stimulus sequences were normalised again.

*Stimulus delivery*

Stimulus delivery was on a computer running Ubuntu 18.04 and Matlab 2018b and used the functions from the psychophysics toolbox version 3 (http://psychtoolbox.org/). The delay between stimulus delivery and triggers sent using a LabJack-U3 (LabJack Corporation, Lakewood, CO, USA) device in the EEG was measured to vary between 0 and 4 ms using a BrainProducts StimTrak (BrainProducts GmbH, Gilching, Germany) device prior to data collection.

*Electrophysiological data collection and reduction*

Scalp electrodes were Ag-AgCl electrodes at the following locations: FP1, FPz, FP2, AF7/AF8 (bipolar), F7, F3, Fz, F4, F8, T7, C3, Cz, C4, T8, P7, P3, Pz, P4, P8, O1, Oz, O2. FCz served as the online reference. Two goldcup electrodes were placed on the mastoids (i.e., A1/A2) for later offline re-referencing. Additionally, a vertical Ag-AgCl EOG electrode was placed below the left eye. Two additional goldcup electrodes were placed as EOG electrodes according to the guidelines of the American Academy of Sleep Medicine and Iber (2007) below the outer canthus of the left and above the outer canthus of the right eye, respectively and referenced online to an additional A1 electrode (Grass® goldcup). Furthermore, we recorded a II-lead ECG as well as EMG activity from two chin electrodes (all Ag-AgCl). During the adaptation night only, participants also wore a thoracic respiration belt, two electrodes to assess leg movements placed on the m. tibialis anterior, and nasal prongs. All Ag-AgCl electrodes belonged to an EasyCap® (Easycap GmbH, Woerthsee-Etterschlag, Germany) EEG cap, which participants wore during most of the time in the lab in the appropriate size. The skin was prepared with abrasive Nuprep® (Weaver and company, Aurora, USA) paste prior to electrode placement. Goldcup electrodes were affixed with Grass® EC2 electrode cream (Natus Medical Inc., Pleasanton, USA), whereas Ag-AgCl electrodes were filled with Elefix® paste (Nihon Koden Europe GmbH, Rosbach, Germany).

The pre-processing pipeline followed the steps previously specified in Blume, del Giudice, Wislowska, Heib, and Schabus (2018). More specifically, data were high-pass filtered at 0.5 Hz. Data acquired during wakefulness but not sleep were corrected for horizontal and vertical eye movement artefacts using an independent-component analysis (ICA), which was performed on all EEG and EOG electrodes. This was followed by manual artefact rejection and low-pass filtering of the data at 30 Hz. If necessary, we manually corrected blink artefacts during nocturnal periods of wakefulness. Finally, the data were referenced to digitally linked mastoids offline and the online-reference FCz was 'regained'. Note that data from both conditions from one participant during sleep were only referenced to A2 as the signal from A1 was noisy throughout both recordings. In six sleep recordings, one electrode was excluded due to technical failure or low signal quality and later interpolated using the Fieldtrip's spline method, in one recording this affected four electrodes. For sleep recordings, we then added information about the sleep stage in which a stimulus was presented as well as the information about the sleep cycle (see below) to the trial information. Subsequently, for the event-related analyses (i.e., event-related potentials [ERPs] and time-frequency analyses), we segmented the data into overlapping segments from -2000 to +2000ms relative to stimulus onset. The segments from the habituation/start phase (i.e., the first 20 trials of a block during wakefulness and the first 10 during sleep) were discarded. For the analyses of EEG slow-wave activity (SWA) in each sleep cycle, continuous data were first segmented in to 30-s epochs and subsequently each 30-s epoch into 15 segments of 2 s. In the latter step, segments containing artefacts were excluded.

For event-related potential (ERP) analyses of the mismatch response, local standard and local deviant trials were baseline-corrected using a 600 ms baseline from -600 to 0 ms relative to the onset of the fifth stimulus. Thus, the baseline spanned the time from the onset of the first to the onset of the last stimulus in a sequence of five. This relatively long baseline was required for the time-frequency analyses. For sleep trials, we additionally distinguished between the four different sleep stages N1, N2,

N3, and REM sleep. We then averaged across local, global, and omission trials within the wakefulness (i.e., evening and morning) and sleep recordings. Note that for the sleep recordings, averaging was performed per sleep stage for each participant. If necessary, the number of trials were balanced, that is, if the available number of trials was lower for local deviants than for local standards, we randomly selected a number of local standard trials to match the number of local deviant trials. For ERP analyses, we considered the time window between the onset of the fifth stimulus (or where it would have been) and 900 ms, which is where the next trial could have started as the minimum ITI was 850 ms.

Besides these event-related potential (ERP) analyses, we also investigated oscillatory responses, that is, event-related synchronisation and desynchronisation (ERS/ERD; Pfurtscheller & Lopes da Silva, 1999). To this end, we calculated time-frequency analyses of individual segments using Morlet complex wavelets ($c$ = 3) on the same trials used for the ERP analyses. Changes in power in a 0-900 ms time window (i.e., 50 ms stimulus + min. 850 ms ITI) relative to stimulus onset were compared to power in a 600 ms baseline. Effects of light exposure were likewise be assessed with cluster-based permutation tests in the 1-15 Hz range.

For the computation of SWA as an indicator of sleep propensity (Dijk & Czeisler, 1995; Borbély, Achermann, Trachsel, & Tobler, 1989; Achermann, Dijk, Brunner, & Borbély, 1993), we calculated SWA across the night within each decile of the NREM part of a sleep cycle. To this end, computed EEG SWA (i.e., delta power between 0.5 and 4.5 Hz) for each decile of each NREM part of a NREM-REM cycle (Feinberg & Floyd, 1979; Rudzik et al., 2018). For the computation of delta power, artefact-free data was segmented into 2-second time bins and subjected to Fast-Fourier transformations (FFT) yielding a frequency resolution of 0.5 Hz. Thereafter, FFT results were averaged in the 0.5-4.5 Hz range within each percentile of each NREM cycle at frontal electrodes F3, Fz, and F4. For each NREM cycle, the analyses thus yielded 10 measures per participant. The analysis procedure described here is based on Chellappa et al. (2013). Note though that the authors had chosen 2-4 Hz as the frequency range of interest for unknown reasons. Finally, the results were exported for statistical evaluation in R (R Core Team, 2015).

For statistical analyses, missing melatonin values (in total 3 values, one of which was in the time window for the statistical analyses) were replaced by the mean between the preceding and the following value. In case this was the overall first (last) sample, the value was replaced by the following (preceding) measurement.

### Supplemental Tables

**Table S1**: Subjective sleep quality reported in the morning of the experimental nights depending on light exposure condition (i.e., mel-low vs. mel-high) and the visit to the lab (i.e., first vs. second visit).

| Effect | b | SE (b) | 95% CI |
|---|---|---|---|
| Condition (mel-low vs. mel-high) | 0.11 | 0.16 | -0.35;0.56 |
| Visit (V1 vs. V2) | -0.06 | 0.23 | -0.51; 0.39 |
| Condition × Visit | 0.14 | 0.38 | -0.62; 0.89 |

Abbreviations: V1/2 = experimental visit 1 vs. 2; b = standardised regression coefficient; SE = Standard Error; CI = Confidence Interval; **p* <.05, ***p* <.01, ****p* <.001.

**Table S2**: Subjective awakening quality reported in the morning of the experimental nights depending on light exposure condition (i.e., mel-low vs. mel-high) and the visit to the lab (i.e., first vs. second visit).

| Effect | b | SE (b) | 95% CI |
|---|---|---|---|
| Condition (mel-low vs. mel-high) | 0.15 | 0.09 | -0.02; 0.31+ |
| Visit (V1 vs. V2) | 0.13 | 0.09 | -0.04; 0.29 |
| Condition × Visit | -0.21 | 0.16 | -0.53; 0.1 |

Abbreviations: V1/2 = experimental visit 1 vs. 2; b = standardised regression coefficient; SE = Standard Error; CI = Confidence Interval; + *p* <.06, **p* <.05, ***p* <.01, ****p* <.001.

**Table S3**: Latency to continuous 10 minutes of sleep, N1, N2, N3, and REM depending on light exposure condition (i.e., mel-low vs. mel-high) and the visit to the lab (i.e., first vs. second visit).

| Latency to | Effect | b | SE (b) | 95% CI |
|---|---|---|---|---|
| Cont. 10 min | Condition (mel-high vs. mel-low) | -9.52 | -1.61 | -2.1; 1.86 |
| | Visit (V1 vs. V2) | -12.2 | -2.2 | -2.44; -1.39 |
| | Condition × Visit | 3.8 | 0.4 | -1.48; 21.92 |
| N1 | Condition (mel-high vs. mel-low) | -2.75 | 6.33 | -15.01; 9.78 |
| | Visit (V1 vs. V2) | -2.15 | 6.33 | -16.48; 8.15 |
| | Condition × Visit | -1.11 | 10.71 | -22.31; 19.85 |
| N2 | Condition (mel-high vs. mel-low) | -11.27 | 5.84 | -22.6; 0.42 |
| | Visit (V1 vs. V2) | -17.04 | 5.84 | -28.6; -5.36 |
| | Condition × Visit | 10.7 | 9.65 | -8.49; 29.58 |
| N3 | Condition (mel-high vs. mel-low) | 2.05 | 6.14 | -9.86; 14.05 |
| | Visit (V1 vs. V2) | -6.69 | 6.14 | -18.87; 5.53 |
| | Condition × Visit | -6.08 | 10.39 | -26.54; 14.47 |
| REM | Condition (mel-high vs. mel-low) | 1.1 | 6.34 | -11.35; 13.45 |
| | Visit (V1 vs. V2) | 5.98 | 6.34 | -6.31; 18.23 |
| | Condition × Visit | -9.6 | 9.61 | -28.1; 9.37 |

Abbreviations: V1/2 = experimental visit 1 vs. 2; b = standardised regression coefficient; SE = Standard Error; CI = Confidence Interval; + *p* <.06, **p* <.05, ***p* <.01, ****p* <.001.

**Table S4**: Percentage of N1, N2, N3, and REM depending on light exposure condition (i.e., mel-low vs. mel-high) and the visit to the lab (i.e., first vs. second visit).

| Percentage of | Effect | b | SE (b) | 95% CI |
|---|---|---|---|---|
| N1 | Condition (mel-high vs. mel-low) | -1.85 | 2.0 | -5.76; 2.09 |
| | Visit (V1 vs. V2) | -2.17 | 2.0 | -6.12; 1.79 |
| | Condition × Visit | 2.17 | 3.79 | -5.32; 9.69 |
| N2 | Condition (mel-high vs. mel-low) | 0.59 | 2.07 | -3.44; 4.7 |
| | Visit (V1 vs. V2) | 0.4 | 2.07 | -3.56; 4.63 |
| | Condition × Visit | -1.12 | 3.69 | -8.53; 6.04 |
| N3 | Condition (mel-high vs. mel-low) | 2.65 | 2.32 | -1.9; 7.28 |
| | Visit (V1 vs. V2) | 2.26 | 2.32 | -2.3; 6.89 |
| | Condition × Visit | -3.83 | 4.35 | -12.45; 4.71 |
| REM | Condition (mel-high vs. mel-low) | -1.38 | 1.81 | -4.98; 2.16 |
| | Visit (V1 vs. V2) | -0.49 | 1.81 | -4.12; 3.07 |
| | Condition × Visit | 2.77 | 3.24 | -3.64; 9.23 |

Abbreviations: V1/2 = experimental visit 1 vs. 2; b = standardised regression coefficient; SE = Standard Error; CI = Confidence Interval; + *p* <.06, **p* <.05, ***p* <.01, ****p* <.001.

**Table S5**: Wake after sleep onset (WASO) depending on light exposure condition (i.e., mel-low vs. mel-high) and the visit to the lab (i.e., first vs. second visit).

| Effect | b | SE (b) | 95% CI |
|---|---|---|---|
| Condition (mel-high vs. mel-low) | 5.96 | 6.11 | -6.07; 17.95 |
| Visit (V1 vs. V2) | 6.13 | 6.11 | -5.96; 17.9 |
| Condition × Visit | -18.99 | 11.01 | -40.29; 2.64 |

Abbreviations: V1/2 = experimental visit 1 vs. 2; b = standardised regression coefficient; SE = Standard Error; CI = Confidence Interval; + *p* <.06, **p* <.05, ***p* <.01, ****p* <.001.

**Table S6**: Sleep efficiency depending on light exposure condition (i.e., mel-low vs. mel-high) and the visit to the lab (i.e., first vs. second visit).

| Effect | b | SE (b) | 95% CI |
|---|---|---|---|
| Condition (mel-high vs. mel-low) | -7.44 | 6.05 | -19.48; 4.81 |
| Visit (V1 vs. V2) | -5.54 | 6.05 | -17.33; 6.52 |
| Condition × Visit | 20.51 | 10.93 | -1.26; 41.56 |

Abbreviations: V1/2 = experimental visit 1 vs. 2; b = standardised regression coefficient; SE = Standard Error; CI = Confidence Interval; + *p* <.06, **p* <.05, ***p* <.01, ****p* <.001.

**Table S7**: Number of awakenings depending on light exposure condition (i.e., mel-low vs. mel-high) and the visit to the lab (i.e., first vs. second visit).

| Effect | b | SE (b) | 95% CI |
|---|---|---|---|
| Condition (mel-high vs. mel-low) | 0.61 | 6.4 | -11.67; 13.09 |
| Visit (V1 vs. V2) | -1.45 | 6.4 | -13.64; 11.1 |
| Condition × Visit | -3.73 | 11.78 | -26.27; 18.83 |

Abbreviations: V1/2 = experimental visit 1 vs. 2; b = standardised regression coefficient; SE = Standard Error; CI = Confidence Interval; + *p* <.06, **p* <.05, ***p* <.01, ****p* <.001.

**Table S8**: Arousal index depending on light exposure condition (i.e., mel-low vs. mel-high) and the visit to the lab (i.e., first vs. second visit).

| Effect | b | SE (b) | 95% CI |
|---|---|---|---|
| Condition (mel-high vs. mel-low) | -0.63 | 4.17 | -8.82; 7.33 |
| Visit (V1 vs. V2) | -1.42 | 4.17 | -9.77; 6.7 |
| Condition × Visit | 0.29 | 8.07 | -15.18; 16.34 |

Abbreviations: V1/2 = experimental visit 1 vs. 2; b = standardised regression coefficient; SE = Standard Error; CI = Confidence Interval; + *p* <.06, **p* <.05, ***p* <.01, ****p* <.001.

**Table S9**: Slow wave activity (0.5-4.5 Hz) depending on light exposure condition (i.e., mel-low vs. mel-high) and the sleep cycle (i.e., first vs. second vs. third cycle).

| **Effect** | **b** | **SE (b)** | **95% CI** |
|---|---|---|---|
| Condition (mel-high vs. mel-low) | 52.45 | 70.21 | -83.38; 191.05 |
| Cycle (1 vs. 2 vs. 3) | -165.37 | 68.83 | -209.01; -119.72 |
| Condition × Cycle | -28.37 | 32.5 | -93.13; 34.76 |

Abbreviations:; b = standardised regression coefficient; SE = Standard Error; CI = Confidence Interval; + *p* <.06, **p* <.05, ***p* <.01, ****p* <.001.

### Supplemental Figures

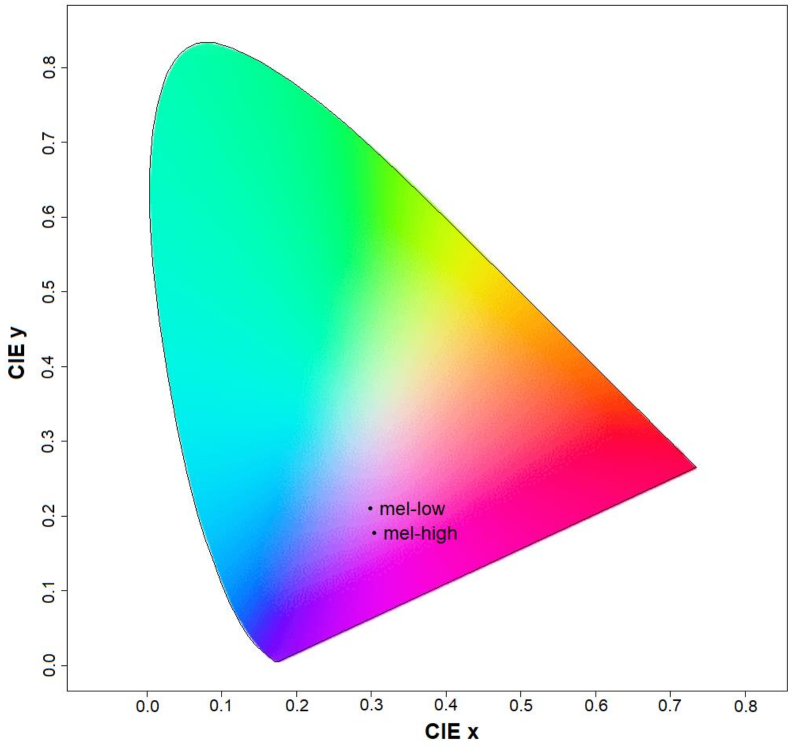


**Suppl. Fig. 1.** Illustration of the irradiance-derived chromaticity coordinates of the two light sources (CIE1931 xy standard observer for a 2° field). Irradiance-derived chromaticity values were x = 0.30 and y = 0.21 for the mel-low and x = 0.30 and y = 0.18 for the mel-high condition. Note that the chromaticities were too far away from the Planckian locus, therefore rendering the correlated colour temperature (CCT), which is defined for colours close to the Planckian locus, unsuitable.

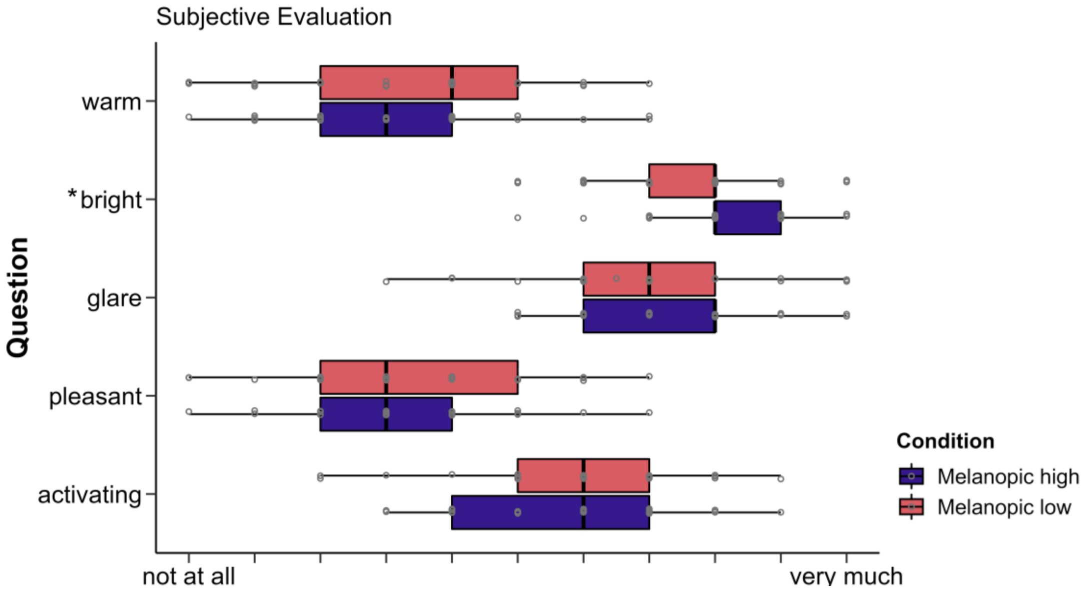


**Suppl. Fig. 2.** Subjective comfort ratings for the two light sources on a 11-item Likert scale regarding warmth, brightness, glare, pleasantness and as how activating volunteers perceived the light. The lower and upper hinges of the boxplot correspond to the 25% and 75% quartiles, the thick black line indicates the median. Whiskers extend to the lowest/largest value at most 1.5× the interquartile range (IQR) from the hinges. The *mel-high* light is perceived as brighter than the *mel-low* light as indicated by the asterisk. There were no differences regarding the other ratings.

### Supplemental Results

#### *Melatonin data*

All individual melatonin curves were inspected visually. One participant turned out to be a high-secretor (15 [high-melanopic] and 9 [low-melanopic] out of 16 samples > 29 pg/ml), wherefore he was excluded from further analyses. In all other participants, an apparent increase in evening melatonin levels was visible, and the light exposure was correctly timed and coincided with the rise in melatonin. Among the remaining 29 participants, 20 presented with lower melatonin values in the *mel-high* compared to the *mel-low* condition 30 min into and just after the light exposure (average suppression 30.2%, range 4.1-77.7%). Two participants showed no difference between the conditions, and in 7 participants, melatonin levels were higher in the *mel-high* condition (average 'suppression' -28.0%, range -0.3 to -97.4%). For an illustration of the individual differences, see Suppl. Fig. 3.

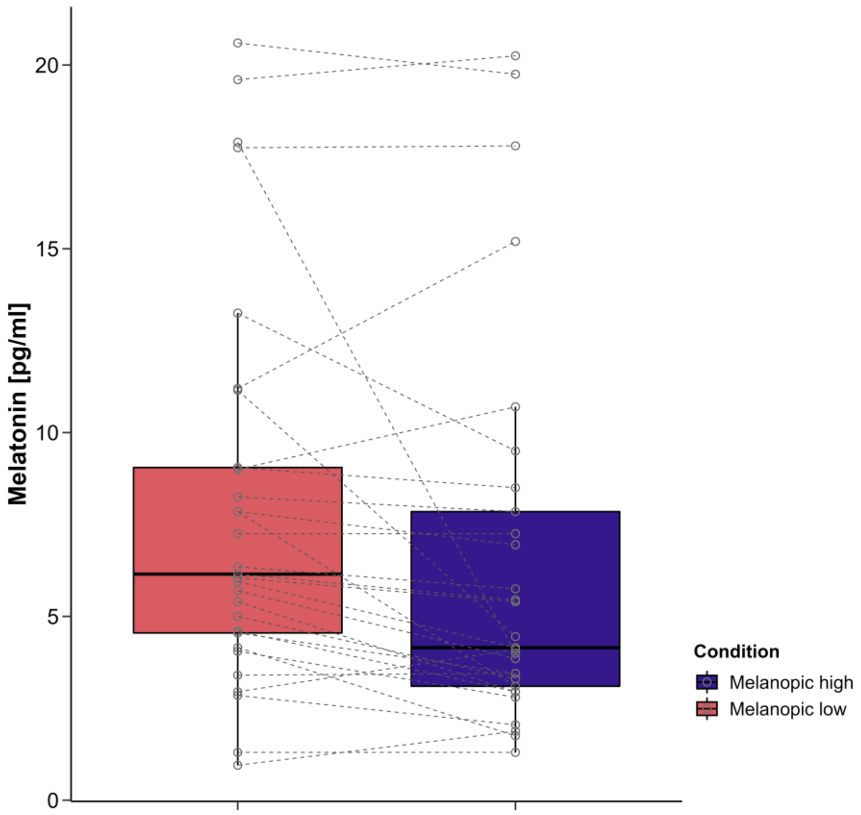


**Suppl. Fig. 3.** Individual differences in salivary melatonin levels between the two light exposure conditions. The data points represent the averaged values 30 min into the light exposure as well as just after the light exposure, i.e., the time points that turned out to differ significantly between the two conditions in the group-level analyses. Of the 29 participants, 20 showed reduced levels in the mel-high condition, 2 had identical values, and 7 had higher levels in the mel-high compared to the mel-low condition.

#### *Subjective Sleepiness (KSS)*

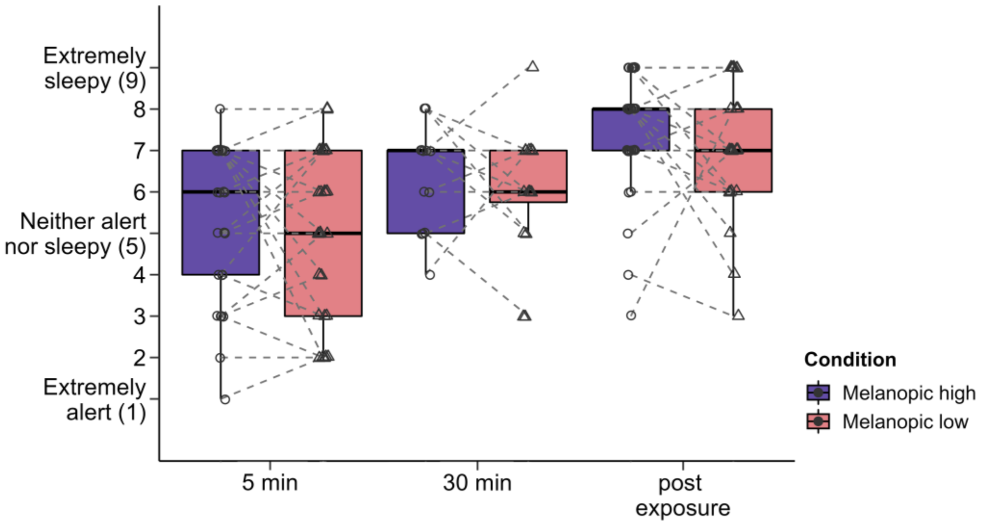


**Suppl. Fig. 4.** Individual differences in the KSS values between the two light exposure conditions 5 minutes into and just after the light exposure. Five minutes into the light exposure, 13 of the 29 participants reported being sleepier in the mel-low compared to the mel-high condition and 10 felt less sleepy while 6 did not report a difference. After 30 min, 5 of 16 respondents (note that this KSS had just been introduced after 13 participants) reported being sleepier in the mel-low compared to the mel-high condition, 8 felt less sleepy, and 3 reported no difference. Just after the light exposure, 7 felt sleepier in the mel-low compared to the mel-high condition, 10 felt less sleepy, and 12 reported no difference.

#### *Area under the curve (AUC)*

In addition to the analyses based on the absolute melatonin values, we also calculated analyses taking into account the AUC. Taking into account all values from the beginning of the light exposure until HBT, the AUC for the melatonin values also differed between the two light conditions. Specifically, the AUC in the *mel-low* condition was larger than in the *mel-high* condition ($F_{ATS}(1)=10.14$, $p=.004$, $RTE_{mel\text{-}low}=0.549$). This was well in line with the analyses of absolute melatonin values. At the two time points, where follow-up analyses of absolute values had revealed significant differences (i.e., 30 minutes in to the light exposure and just after the exposure), the AUC was smaller by 14.6 % in the *mel-high* relative to the *mel-low* condition. Similarly to the results obtained using absolute values, there were no differences between the AUCs for the melatonin values prior to the start of the light exposure ($F_{ATS}(1)=0.77$, $p=.57$, $RTE_{mel\text{-}low}=0.518$) nor the values in the morning ($F_{ATS}(1)=0.27$, $p=.6$, $RTE_{mel\text{-}low}=0.49$).

***Time-frequency analyses***

*Wakefulness*

During wakefulness, there was also a significant mismatch effect with deviants resulting in stronger ERS (mel-low: $p < .001$; mel-high: $p < .001$ & $p = .032$; Suppl. Fig. 4 A/F). However, there were no differences between the low- and high-melanopic light exposure conditions (all $p > .12$).

*Sleep*

*N1*

During N1, there was a continued mismatch response (mel-low: $p < .001$; mel-high: $p < .001$; Suppl. Fig. 4 B/G) suggesting a continued mismatch response. There were no differences between the light exposure conditions (all $p > .24$).

*N2*

In N2, the mismatch effect persisted (mel-low: $p < .001$; mel-high: $p < .001$; Suppl. Fig. 4 C/H). There were no differences between the two light exposure conditions (all $p > .27$).

*N3*

Even in N3, the mismatch effect persisted (mel-low: $p < .001$; mel-high: $p < .001$; Suppl. Fig. 4 D/I). There was a significant difference between the two light exposure conditions, where the mel-low condition elicited slightly stronger effects than the mel-high condition (at electrode Cz: between 0 and 500 ms; $p = .039$).

*REM*

During REM, there were significant mismatch effects (mel-low: $p < .001$; mel-high: $p < .001$; Suppl. Fig. 4 E/J), which were limited to earlier time windows and not as strong as during NREM and wakefulness. There were no differences between the two light-exposure conditions (all $p > .61$).

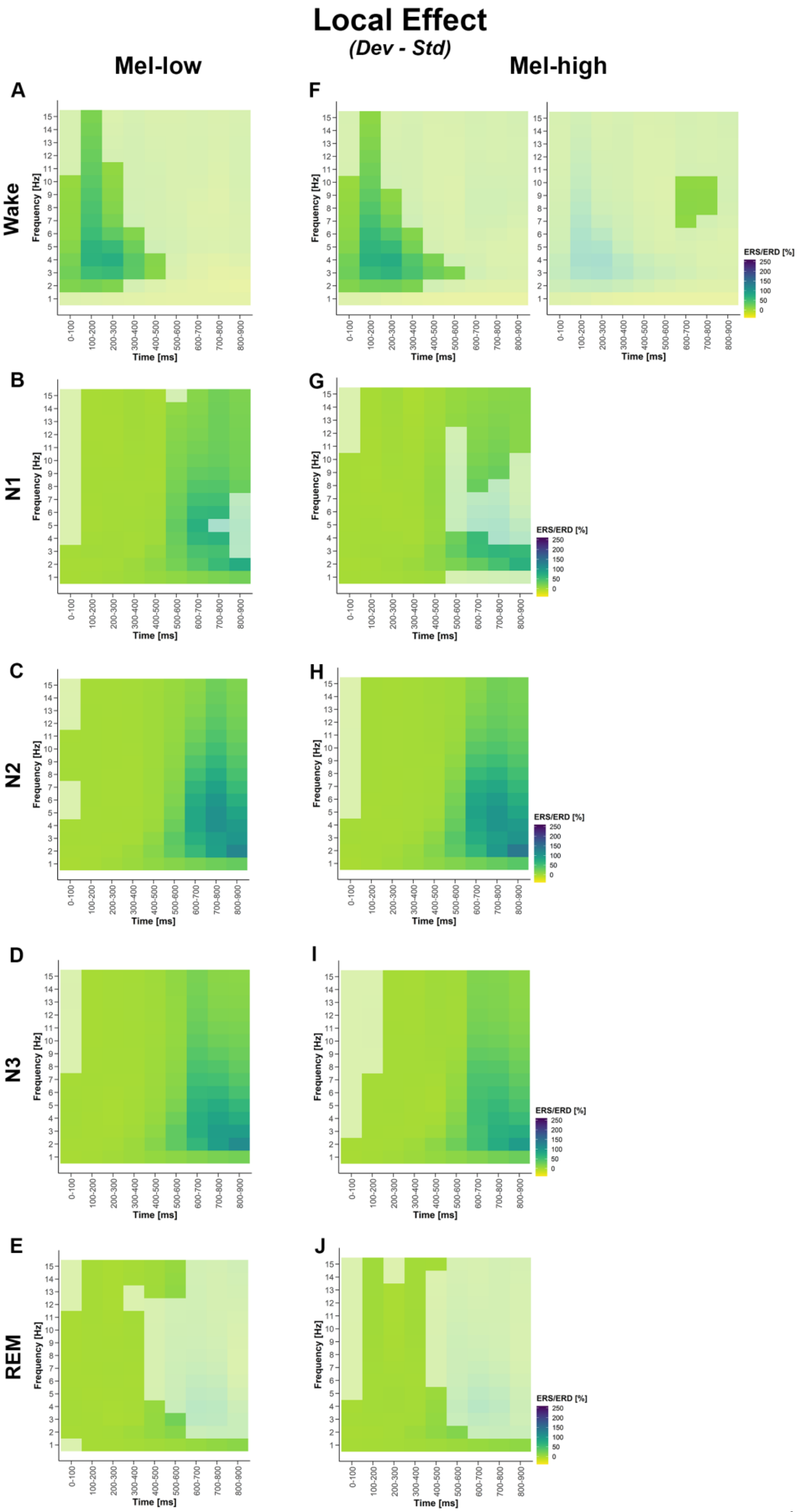
Local Effect
(Dev - Std)
Mel-low
Mel-high
A
F
B
G
C
H
D
I
E
J
Wake
N1
N2
N3
REM
Frequency [Hz]
Time [ms]
0-100
100-200
200-300
300-400
400-500
500-600
600-700
700-800
800-900
ERS/ERD [%]
250
200
150
100
50
0

**Suppl. Fig. 5.** Mismatch effects from time-frequency analyses between 1-15 Hz during wakefulness (A, F), N1 (B, G), N2 (C, H), N3 (D, I), and REM (E, J) in the mel-high and the mel-low conditions. The plots show the difference between deviants and standards at electrode Cz across frequencies and time windows (0-900 ms in 100 ms windows). Rich colours indicate significant clusters. Event-related synchronization (ERS) corresponds to deviants eliciting a stronger increase in power compared to standards. Only significant clusters are shown (cf. main supplementary text for more information).

### Supplemental Lighting Information

The photon density between 440 to 540 nm, that is, the full width at half maximum (FWHM) for melanopsin, was $2.16 \times 10^{13} \frac{photons}{\sec \times cm^2}$ in the *mel-low* and $4.65 \times 10^{13} \frac{photons}{\sec \times cm^2}$ in the *mel-high* condition. According to the following formula, the photon density per second and square centimetre can be calculated from the provided spectral data (see below) for any range of wavelengths: $N_p = \sum_{k=low}^{high} I \times \lambda_k \times 5.03 \times 10^{11}$, where $I$ is the irradiance measure and $\lambda$ is the wavelength in nm.

We additionally provide a CSV-file ("BLUMEC_spectral_ir-radiance_tab.csv") specifying spectral (ir-)radiances for each wavelength between 380 and 780 nm. Spectral *irradiance* (W/[sqm*nm], columns 2 and 3) and *radiance* (W/[sr*m$^2$], columns 4 and 5) measures of the experimental conditions were taken at a distance of 70 cm from the screen at a height of 120 cm, that is, from the observer's point of view. Horizontal *radiance* was measured at a distance of 20 cm from the screen at the centre of the screen (C, columns 9 and 12), 20 cm to the left (L, columns 10 and 13), and to the right (R, columns 11 and 14). Additionally, we provide the spectral radiance for the background room light for the evening room light (column 6), the evening dim light condition (column 7), and the morning room light (column 8). For the measurements of the background light, the measuring device was turned in the direction of the light source from the position of the participants' heads. We recommend the users to use the luox app (www.luox.app) to calculate the radiance-derived α-opic responses ($L_{e,\Omega}$; in mW/m$^2$*sr), radiance-derived α-opic equivalent daylight (D65) luminances (EDL, in cd/m$^2$), irradiance-derived α-opic responses ($E_e$; in mW/m$^2$), and equivalent daylight (D65) illuminances from the data.

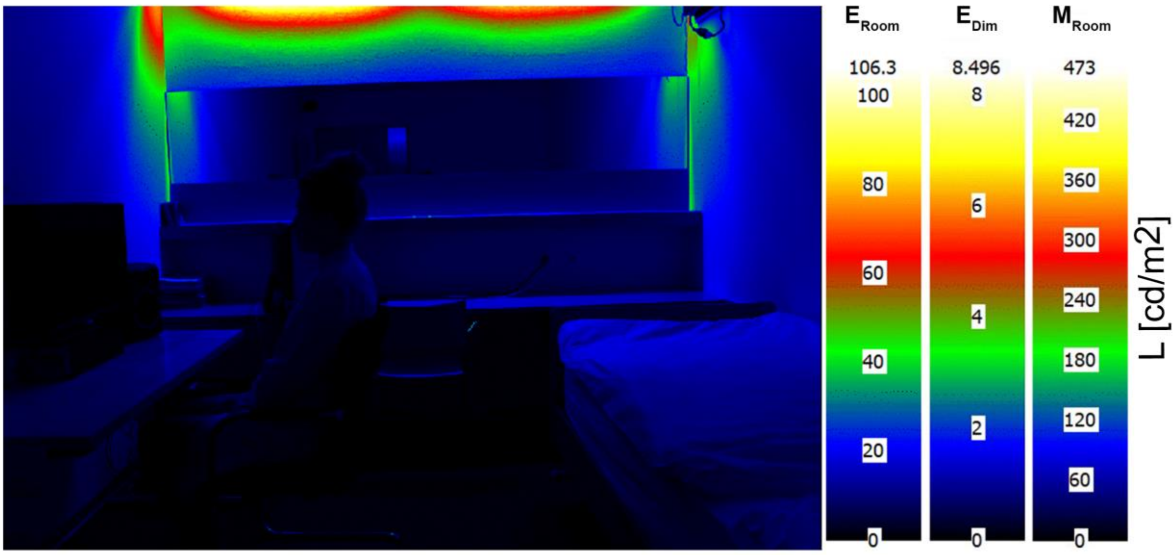


**Suppl. Fig. 6.** Luminance densities in cd/m$^2$ for the background lighting, i.e., "evening room" ($E_{Room}$), "evening dim" ($E_{Dim}$), and "morning room" ($M_{Room}$).
